# Tick hemocytes have pleiotropic roles in microbial infection and arthropod fitness

**DOI:** 10.1101/2023.08.31.555785

**Authors:** Agustin Rolandelli, Hanna J. Laukaitis-Yousey, Haikel N. Bogale, Nisha Singh, Sourabh Samaddar, Anya J. O’Neal, Camila R. Ferraz, Matthew Butnaru, Enzo Mameli, Baolong Xia, M. Tays Mendes, L. Rainer Butler, Liron Marnin, Francy E. Cabrera Paz, Luisa M. Valencia, Vipin S. Rana, Ciaran Skerry, Utpal Pal, Stephanie E. Mohr, Norbert Perrimon, David Serre, Joao H.F. Pedra

**Author notes:** Present address: Rancho BioSciences, San Diego, CA 92127. Present address: Immunology Program, Memorial Sloan Kettering Cancer Center, New York, NY 10065.

## Abstract

Uncovering the complexity of systems in non-model organisms is critical for understanding arthropod immunology. Prior efforts have mostly focused on Dipteran insects, which only account for a subset of existing arthropod species in nature. Here, we describe immune cells or hemocytes from the clinically relevant tick *Ixodes scapularis* using bulk and single cell RNA sequencing combined with depletion via clodronate liposomes, RNA interference, Clustered Regularly Interspaced Short Palindromic Repeats activation (CRISPRa) and RNA-fluorescence *in situ* hybridization (FISH). We observe molecular alterations in hemocytes upon tick infestation of mammals and infection with either the Lyme disease spirochete *Borrelia burgdorferi* or the rickettsial agent *Anaplasma phagocytophilum*. We predict distinct hemocyte lineages and reveal clusters exhibiting defined signatures for immunity, metabolism, and proliferation during hematophagy. Furthermore, we perform a mechanistic characterization of two *I. scapularis* hemocyte markers: *hemocytin* and *astakine*. Depletion of phagocytic hemocytes affects *hemocytin* and *astakine* levels, which impacts blood feeding and molting behavior of ticks. Hemocytin specifically affects the c-Jun N-terminal kinase (JNK) signaling pathway, whereas astakine alters hemocyte proliferation in *I. scapularis*. Altogether, we uncover the heterogeneity and pleiotropic roles of hemocytes in ticks and provide a valuable resource for comparative biology in arthropods.

## Introduction

The evolution of arthropod immune systems is shaped by the environment, longevity, nutrition, microbial exposure, development, and reproduction (*1–3*). Yet, our current understanding of immunology is centered around Dipteran insects, which reflects an incomplete depiction of arthropod biological networks existing in nature (*4–10*). Divergent signaling modules prevail between Dipterans and other arthropods, highlighting the complex and dynamic features occurring across evolutionarily distant species. One example is the immune deficiency (IMD) pathway, a network analogous to the tumor necrosis factor receptor (TNFR) in mammals (*11, 12*). Some components of the canonical IMD pathway are not observed in arachnids, although this immune pathway remains functional and responsive to microbial infection (*4–8*). Another instance is the janus kinase/signal transducer and activator of transcription (JAK/STAT) signaling pathway, which in flies is elicited by unpaired cytokine-like molecules (*13*), whereas in ticks host-derived interferon (IFN)-g facilitates Dome1-mediated activation (*9, 10*). This biochemical network enhances tick blood meal acquisition and development while inducing the expression of antimicrobial components (*9, 10*). Thus, investigating the plasticity of immune networks in evolutionarily divergent organisms may reveal discrete aspects of arthropod biology.

Ticks are ancient arthropods that serve as major vectors of human and animal pathogens, including the Lyme disease spirochete *Borrelia burgdorferi* and the rickettsial agent *Anaplasma phagocytophilum* (*14–16*). Selective pressure caused by longevity (*i.e.*, two to ten years in certain species), exclusive hematophagy in all life stages, long-term dispersal (adaptation to multiple environments) and exposure to an array of microbes, make ticks distinct from other hematophagous arthropods (*17, 18*). While it is acknowledged that the immune system plays an important role in vector competence, our understanding of the mechanisms by which immune resistance balances microbial infection in ticks remain fragmented (*6–10*).

Hemocytes are specialized arthropod immune cells that function in both cellular and humoral responses. These cells circulate through the hemolymph and are in contact with tissues within the arthropod body cavity (*19*). Historically, tick hemocytes have been categorized according to their cellular morphology and ultrastructural characteristics (*20, 21*). Although useful, this classification is incomplete because the ontogeny, plasticity, and molecular features of hemocytes during hematophagy and infection remain obscure. Here, we used bulk and single cell RNA sequencing (scRNA-seq) coupled with Clustered Regularly Interspaced Short Palindromic Repeats (CRISPR) activation, RNA interference (RNAi), RNA-fluorescence *in situ* hybridization (RNA-FISH), and immune cell depletion through treatment with clodronate liposomes to reveal distinct features of hemocyte clusters in the blacklegged tick. We identify marker genes for each hemocyte cluster, predict lineages, and unveil specific biological signatures related to immunity, proliferation, and metabolism during hematophagy. We further characterize an immune cluster that expresses *hemocytin* (*hmc*) and *astakine* (*astk*).

Manipulation of phagocytic hemocytes and the expression of *hmc* or *astk* impacted bacterial acquisition, feeding and molting. Overall, we highlight the exquisite biology of ticks, demonstrate the canonical and non-canonical roles of immune cells in an obligate hematophagous ectoparasite, and provide a critical resource for comparative biology of arthropods.

## Results

### Blood feeding induces molecular signatures in *I. scapularis* hemocytes related to immunity, metabolism, and proliferation

Ticks rely solely on blood as a source of essential metabolites, ingesting ∼100 times their body mass per meal (*22*). During feeding, extensive modifications and tissue rearrangements are necessary to accommodate and digest such large volumes of blood (*22*). Given their strategic position as circulating cells in the hemolymph, we hypothesized that hemocytes sense and respond to physiological and microbial exposure during tick infestation on mammals. We optimized hemocyte isolation from *I. scapularis* nymphs, the clinically relevant stage in the blacklegged tick, and identified three common hemocyte morphotypes reported in the literature (Figures 1A-B) (*20, 21*). Prohemocytes, considered the stem cell-like hemocyte population, were the smallest cells, with a round or oval shape of ∼5-10 µm. The cytoplasm area was minute (high nuclear/cytoplasmic ratio) and homogeneous, with no apparent protrusions or granules (Figure 1B). Plasmatocytes varied in size (∼15-30 µm) and shape (oval, ameboid-like, pyriform), and had cytoplasmic protrusions or pseudopodia-like structures. The cytoplasm was clear and had few dark-blue or violet stained granules or vacuoles. The nucleus was located near the center of the cell (Figure 1B). Granulocytes were ∼10-20 µm round or oval shape cells. The position of the nucleus varied, appearing most often near the periphery of the cell. Their cytoplasm was filled with dark-blue or violet stained granules or vacuoles (Figure 1B). We then investigated the impact of blood-feeding on hemocytes originating from *I. scapularis* nymphs. We observed an increased quantity of total hemocytes upon mammalian feeding (Figure 1C). The percentage of plasmatocytes increased in engorged ticks, whereas we noticed a decline in the proportion of prohemocytes and granulocytes from unfed compared to repleted nymphs (Figure 1C). Altogether, we demonstrate the impact of hematophagy on the distribution of tick hemocyte morphotypes.

**Figure 1:**
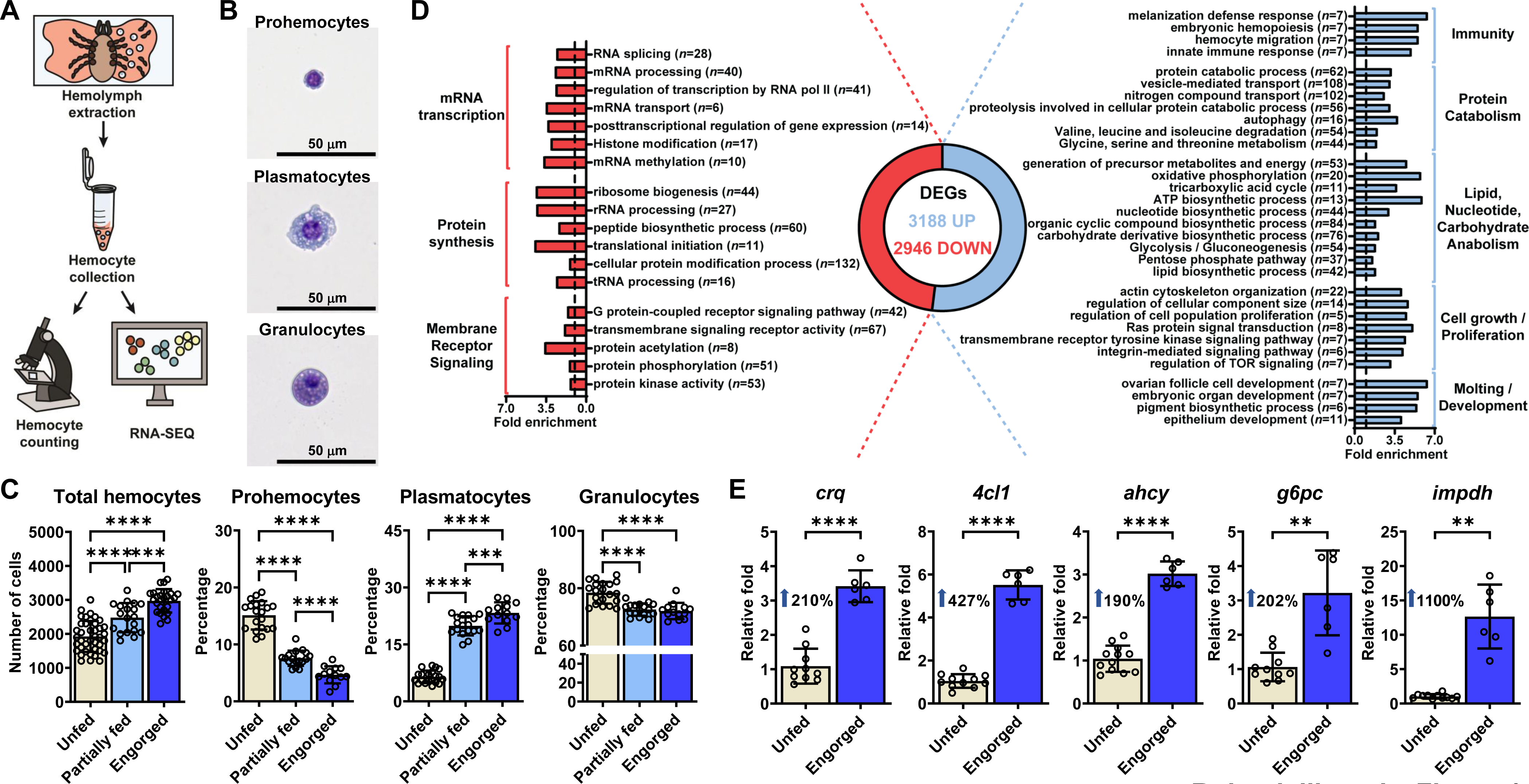
Blood-feeding induces alterations in *I. scapularis* hemocytes. **(A)** Schematic representation of the hemocyte collection procedure. **(B)** Three main morphological subtypes of *I. scapularis* hemocytes evaluated by bright field microscopy after staining. **(C)** Total number of hemocytes and morphotype percentages from unfed (ivory), partially fed (light blue) or engorged (dark blue) nymphs (*n*=13-40). **(D)** Functional enrichment analysis of the differentially expressed genes (DEGs) present in hemocytes from engorged ticks (blue; Up) compared to unfed ticks (red; Down). Fold enrichment of significant terms are depicted. Number of DEGs per category are shown in parentheses. **(E)** The expression of immune, metabolic and proliferative genes in hemocytes from unfed (ivory) and engorged (dark blue) ticks was evaluated by RT-qPCR (*n*=6- 12; samples represent 40-80 pooled ticks). Results are represented as mean ± SD. At least three biological replicates were performed. Statistical significance was evaluated by **(C)** ANOVA or **(E)** an unpaired t-test with Welch’s correction. \*\**p*<0.01; \*\*\*\**p*<0.0001. *crq* = croquemort; *4cl1* = 4-coumarate-CoA ligase 1; *ahcy* = adenosylhomocysteinase B; *g6pc* = glucose-6- phosphatase 2; *impdh* = inosine-5’-monophosphate dehydrogenase 1.

Next, we aimed to examine global transcriptional changes induced by hematophagy through bulk RNA-seq. We collected hemocytes from unfed and engorged nymphs and observed drastic changes in gene expression with a total of 6,134 differentially expressed genes (DEGs) (Figure 1D; Supplementary Dataset 1). Hemocytes collected from unfed *I. scapularis* nymphs were enriched for housekeeping genes, such as mRNA transcription (*e.g.*, RNA splicing, mRNA processing, histone modification), protein synthesis (*e.g.*, peptide biosynthetic process, cellular protein modification process, ribosome biogenesis) and membrane receptor signaling pathways (*e.g.*, transmembrane signaling receptor activity, G protein-coupled receptor signaling pathway, protein kinase activity) (Figure 1D; Supplementary Figure 1; Supplementary Dataset 2).

Conversely, hemocytes originating from engorged ticks were enriched in gene signatures related to immunity, metabolic pathways, cell proliferation/growth, and arthropod molting/development (Figure 1D; Supplementary Figure 2; Supplementary Dataset 2). Key genes upregulated during feeding in hemocytes were independently validated through the detection of *croquemort* (*crq*), *4-coumarate-CoA ligase 1* (*4cl1*), *adenosylhomocysteinase B* (*ahcy*), *glucose-6-phosphatase 2* (*g6pc*), and *inosine-5’-monophosphate dehydrogenase 1* (*impdh*) via qRT-PCR (Figure 1E). Overall, the data indicate that *I. scapularis* hemocytes express a dynamic genetic program during hematophagy.

### Defining *I. scapularis* hemocyte clusters by scRNA-seq

To uncover whether the transcriptional changes observed through bulk RNA-seq is accompanied with heterogeneity among hemocytes, we performed scRNA-seq. We collected hemocytes from (1) unfed nymphs, (2) engorged nymphs fed on uninfected mice, and engorged nymphs fed on mice infected with either (3) the rickettsial pathogen *A. phagocytophilum* or (4) the Lyme disease spirochete *B. burgdorferi*. After stringent quality controls, we profiled a total of 20,432 cells (unfed = 4,630; engorged uninfected = 6,000; engorged *A. phagocytophilum*-infected = 6,287; engorged *B. burgdorferi*-infected = 3,515), with a median of 744 unique molecular identifiers (UMIs), 261 genes and 6.2% of mitochondrial transcripts per cell across conditions (Supplementary Figure 3). Consistent with the bulk RNA-seq results, the principal component analysis (PCA) identified distinct distributions between unfed and fed conditions, reinforcing the notion that significant cellular and/or transcriptional changes occur following a blood meal (Supplementary Figure 4).

Following unsupervised clustering, we identified seven clusters in unfed ticks and thirteen clusters in engorged nymphs (Supplementary Figure 5). Based on similarities in marker gene profiles, two clusters in the unfed and two clusters in the engorged datasets were merged (Supplementary Datasets 3-4). Thus, six and twelve clusters remained, respectively, with each cluster expressing a unique set of cell type-defining genes (Figure 2A-C, Supplementary Datasets 5-6). One cluster in each dataset showed high expression of gut-associated genes *cathepsinB*, *cathepsinD* and *boophilinH2* (Figure 2A-C, Supplementary Dataset 5-6) and two additional clusters in the engorged dataset had gene expression profiles indicative of cuticle and salivary glands tissues (Figure 2A-C, Supplementary Dataset 6). Therefore, these clusters were excluded from subsequent analysis. We assigned putative functions to the top 50 DEGs per cluster using information for tick genes in VectorBase, the sequence homology to *D. melanogaster* in FlyBase and the presence of functionally annotated protein domains in InterPro (Figure 2C-E, Supplementary Dataset 5-6). We also performed functional enrichment analysis based on gene ontology (GO) using the entire list of DEGs for each cluster (Supplementary Dataset 7-8). Altogether, we defined five hemocyte clusters from unfed and nine clusters in engorged *I. scapularis* nymphs.

**Figure 2:**
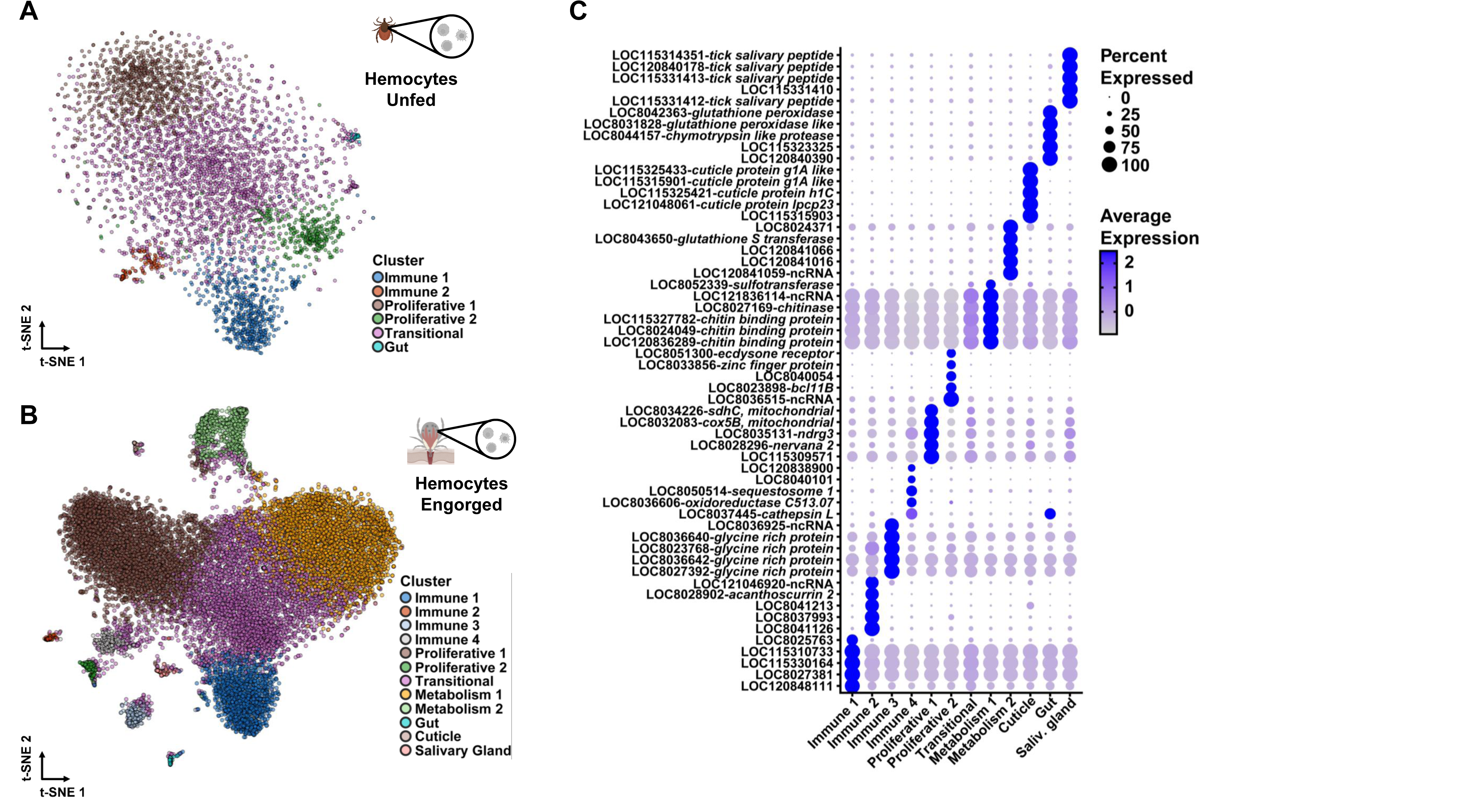
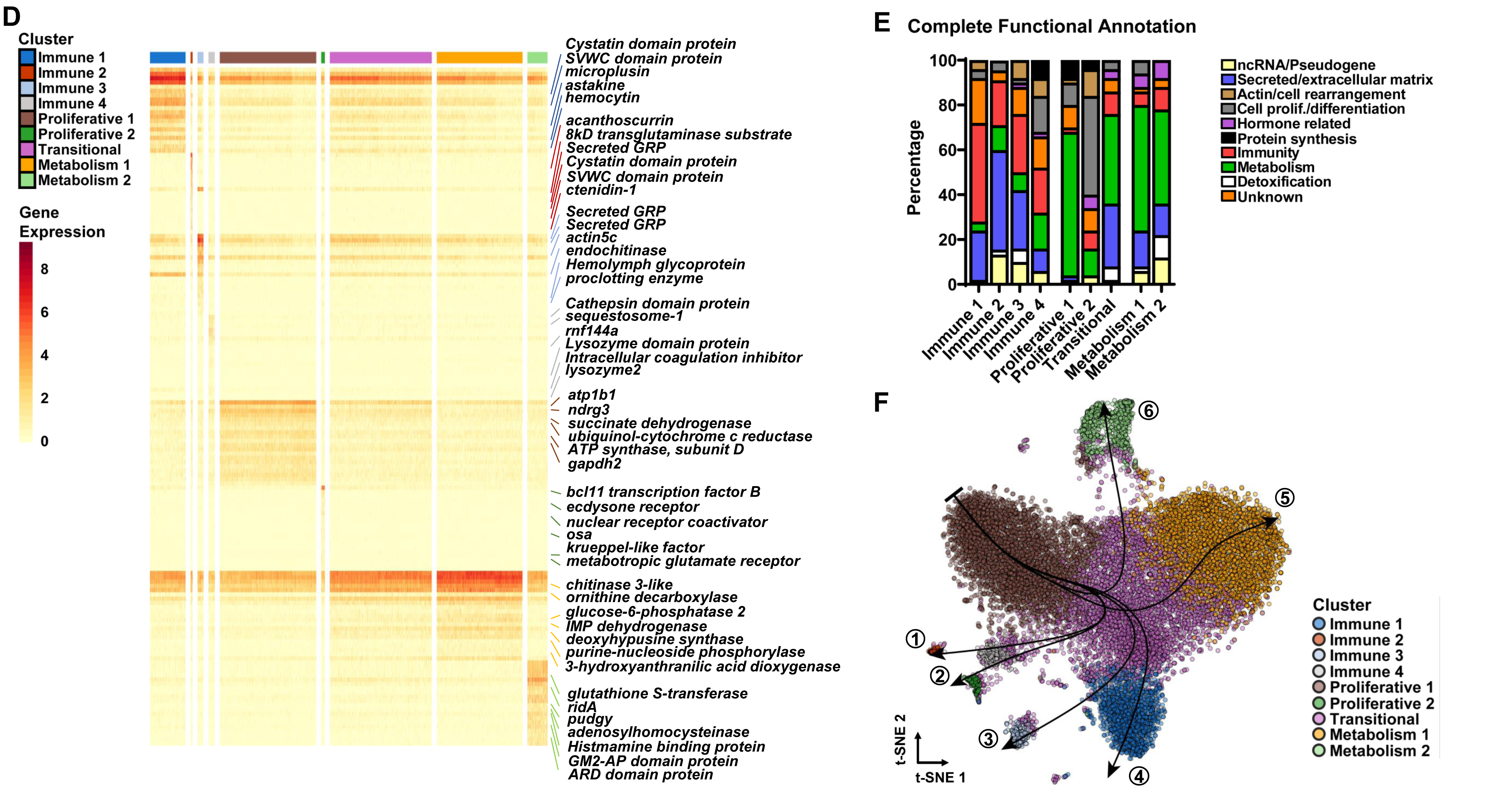
scRNA-seq uncovers hemocytes with immune, proliferative and metabolic signatures in *I. scapularis*. t-Distributed Stochastic Neighbor Embedding (t-SNE) plot clustering of cells collected from the hemolymph of **(A)** unfed (4,630 cells) and **(B)** engorged (15,802 cells) *I. scapularis* nymphs. The engorged t-SNE contains cells from uninfected (6,000 cells), *A. phagocytophilum*-infected (6,287 cells) and *B. burgdorferi*-infected (3,515 cells) *I. scapularis*. **(C)** Dot plot of the top 5 marker genes present in clusters from engorged ticks. Average gene expression is demarked by intensity of color. Percent of gene expression within individual clusters is represented by the dot diameter. **(D)** Heatmap depicting expression of the top 20 marker genes present in hemocyte subtypes from engorged ticks. Representative genes per cluster are highlighted. **(E)** The top 50 marker genes from each hemocyte cluster were manually annotated using publicly available databases, such as VectorBase, FlyBase, and UniProt. The percentage of predicted functional categories, such as ncRNA/pseudogenes (yellow), protein synthesis (black), secreted/extracellular matrix (blue), unknown (orange), actin/cell rearrangement (brown), detoxification (white), cell proliferation/differentiation (grey), metabolism (green), hormone-related (purple), and immunity (red) are shown. **(F)** Pseudotime analysis defined six hemocyte lineages (arrows) in engorged ticks.

### scRNA-seq uncovers hemocyte adaptations to hematophagy in *I. scapularis*

The molecular features and differentiation process of tick hemocytes is currently unknown. Thus, based on GO enrichment and marker gene profiles, we were able to characterize clusters of hemocytes shared by both unfed and engorged ticks (Supplementary Figure 6-7, Supplementary Dataset 5-6). The Immune 1 cluster showed high expression of genes related to phagocytosis or cytoskeleton organization; coagulation and agglutination functions, such as lectins (*hemocytin*, *techylectin-5A*), chitin-binding and clotting proteins; and secreted proteins related to immunity, such as *astakine*, *microplusin*, mucins and cystatin domain peptides (Figure 2C-E, Supplementary Dataset 5-6). The Immune 2 cluster displayed genes encoding secreted proteins involved in immunity, such as antimicrobial peptides (AMPs) and clotting related peptides (Figure 2C-E, Supplementary Dataset 5-6). The Proliferative 1 cluster was enriched with mitochondrial genes, characteristic of stem cells in ancient arthropods, such as crayfish (*23*). This cell cluster also had high expression of genes related to actin polymerization, cell proliferation and differentiation (Figure 2C-E, Supplementary Dataset 5-6). The Proliferative 2 cluster displayed a high percentage of transcription factors, RNA binding proteins and genes related to actin dynamics. Marker genes for this cluster included several genes involved in hormone-related responses, suggesting they may be responsive to ecdysteroids synthesized after a blood meal (Figure 2C-E, Supplementary Dataset 5-6). Lastly, both datasets had a Transitional cluster indicative of intermediate subtypes (Figure 2C-E, Supplementary Dataset 5- 6).

Four hemocyte clusters were only observed in engorged *I. scapularis* ticks. The Immune 3 cluster displayed an enrichment in secreted proteins and genes related to immune functions. This cluster was enriched for chitinases, matrix and zinc metalloproteinases, peptidases and actin binding proteins, which have roles related to wound healing or tissue rearrangement.

Several glycine-rich proteins (GRPs), commonly associated with antimicrobial properties or structural proteins, were also present (Figure 2C-E, Supplementary Dataset 5-6). The Immune 4 cluster showed an overrepresentation of genes related to protein degradation, immune function and cell proliferation (Figure 2C-E, Supplementary Dataset 5-6). Thus, we posit that the Immune 4 cluster represents an intermediate state between the Immune 2 and the Proliferative 2 clusters. Two clusters displayed an enrichment for genes related to metabolic functions. The Metabolism 1 cluster represented genes involved in sulfonation of proteins, lipids and glycosaminoglycans, transmembrane solute transporter, nucleotide and protein metabolism (Figure 2C-E, Supplementary Dataset 5-6). The Metabolism 2 cluster displayed genes related to detoxification, histamine binding, lipid metabolism, methionine metabolism and synthesis of juvenile hormone (Figure 2C-E, Supplementary Dataset 5-6).

Based on these findings, we predicted hemocyte ontogeny using pseudotime analysis (Figure 2F). We found six trajectories considering the cluster Proliferative 1 as a stem cell-like subpopulation. Lineage 1 and 2 ended with the Immune 2 and Proliferative 2 clusters, respectively, with the Immune 4 cluster serving as an intermediate state. Lineages 3 and 4 gave rise to the Immune 3 and Immune 1 clusters, respectively. Finally, two lineage trajectories ended with the metabolic clusters, Metabolism 1 and Metabolism 2. Overall, our findings suggest the presence of an oligopotent subpopulation that differentiates into more specialized subtypes involved in immune and metabolic functions, a process evoked by hematophagy.

### Bacterial infection of *I. scapularis* ticks alters the molecular profile of hemocytes

The impact of bacterial infection on subtypes of tick hemocytes remains elusive. Thus, we collected hemocytes from *I. scapularis* nymphs fed on uninfected, *A. phagocytophilum-* or *B. burgdorferi*- infected mice and determined morphotype percentages. During *A. phagocytophilum* infection, a relative decrease in prohemocytes and increase in plasmatocytes was noted (Figure 3A). Only a slight decrease in the proportion of prohemocytes was observed during *B. burgdorferi* infection of ticks (Figure 3A). No difference in total hemocyte numbers was observed across infection conditions (Supplementary Figure 8). However, partitioning the engorged scRNA-seq datasets by treatments revealed a reduction in the Transitional cluster with an expansion in the Metabolism 1 cluster during *B. burgdorferi* infection (Figure 3B-C). We next analyzed transcriptional changes at the cellular level in engorged uninfected nymphs and compared to engorged ticks infected with the rickettsial agent *A. phagocytophilum* or the Lyme disease spirochete *B. burgdorferi*. Hemocyte clusters were grouped according to three molecular programs: “Immune” (Immune 1-4), “Proliferative” (Proliferative 1-2 and Transitional) and “Metabolism” (Metabolism 1-2). We found 13 DEGs within Immune clusters, 322 DEGs among Proliferative clusters, and 109 DEGs between Metabolism clusters during *A. phagocytophilum* or *B. burgdorferi* infection (Figure 3D, Supplementary Dataset 9). Moreover, 42% (5 out of 12) of genes differentially expressed across all subtypes were marker genes of the Immune 1 cluster (Figure 3D). Collectively, both *A. phagocytophilum* and *B. burgdorferi* affect the molecular signatures of hemocytes in fed *I. scapularis* nymphs.

**Figure 3:**
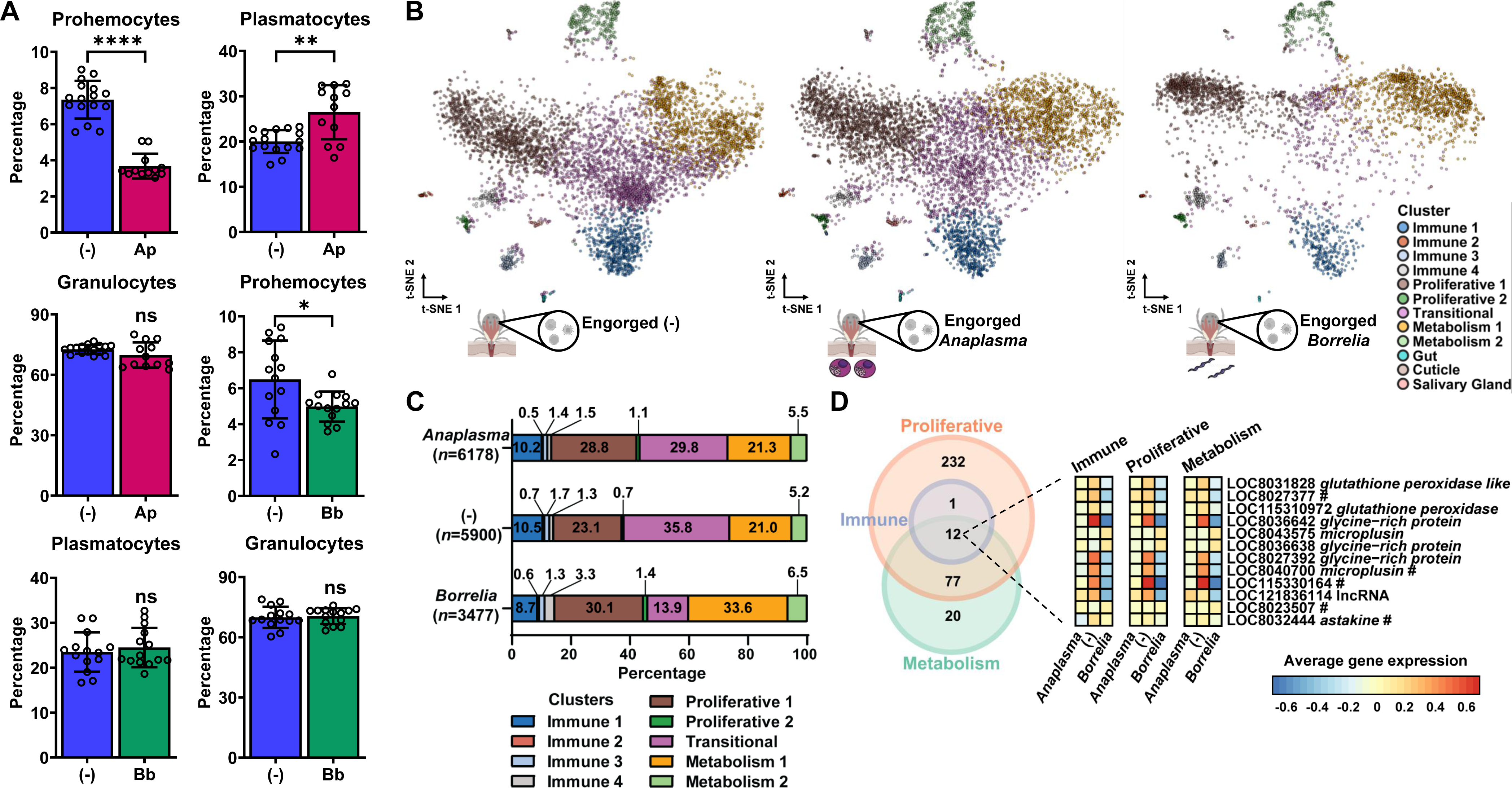
Bacterial infection alters hemocyte subtypes and their molecular expression. **(A)** Hemocyte morphotypes in *I. scapularis* nymphs fed on *A. phagocytophilum-* (Ap, pink) or *B. burgdorferi*- (Bb, green) infected mice compared to uninfected [(-), dark blue] (*n*=10-16). Results are represented as mean ± SD. At least two biological replicates were performed. Statistical significance was evaluated by an unpaired t-test with Welch’s correction. \**p* < 0.05; \*\**p* < 0.01; \*\*\*\**p*<0.0001. ns= not significant. **(B)** t-Distributed Stochastic Neighbor Embedding (t-SNE) plot clustering of cells collected from the hemolymph of uninfected (6,000 cells), *A. phagocytophilum-* (6,287 cells) or *B. burgdorferi*-infected (3,515 cells) *I. scapularis* nymphs. **(C)** Percent of hemocyte clusters present in each experimental condition. **(D)** Venn diagram (left) depicting the number of differentially expressed genes (DEGs) during infection between clusters grouped by putative function (immune, proliferative or metabolic). Heatmap (right) representing the change in expression patterns of DEGs during infection shared between all 3 cluster groups. DEGs were determined using pairwise comparisons against uninfected. # = Immune 1 marker gene. (-) = Uninfected. *Anaplasma* = *A. phagocytophilum*. *Borrelia* = *B. burgdorferi*.

### Hemocytin and astakine contribute to *A. phagocytophilum* infection in *I. scapularis*

The findings described thus far suggested that the Immune 1 hemocyte cluster was an important subpopulation of cells affected by bacterial infection of ticks (Figure 3D). Therefore, we next focused on two marker genes from the Immune 1 hemocyte cluster: *hemocytin* and *astakine* (Figure 4A, Supplementary Dataset 5-6). *Hemocytin* is homologous to *hemolectin* in *D. melanogaster* and von Willebrand factors of mammals (*24–26*). *Hemocytin* encodes for a large multidomain adhesive protein involved in clotting, microbial agglutination and hemocyte aggregation (*24–26*). Conversely, astakine is a cytokine-like molecule present in chelicerates and crustaceans and is homologous to vertebrate prokineticins (*27–30*). Astakine also induces hemocyte proliferation and differentiation of immune cells (*27–30*). *Hemocytin* and *astakine* were broadly expressed in the Immune 1 cluster of *I. scapularis* hemocytes (Figure 4A). We confirmed the expression of *hemocytin* and *astakine* in fixed hemocytes collected from unfed ticks through RNA-FISH (Figure 4B). Like the scRNA-seq profile, a higher percentage of hemocytes exhibited dual expression of both markers, whereas a few cells expressed only one gene. Furthermore, we noted an upregulation of *hemocytin* and *astakine* in hemocytes collected from engorged ticks, which was also observed for other markers of the Immune 1 cluster by bulk RNA-seq (Figure 4C, Supplementary Dataset 10). These findings suggest that blood feeding expands this hemocyte subtype and/or upregulates the expression of its marker genes in *I. scapularis* ticks.

**Figure 4:**
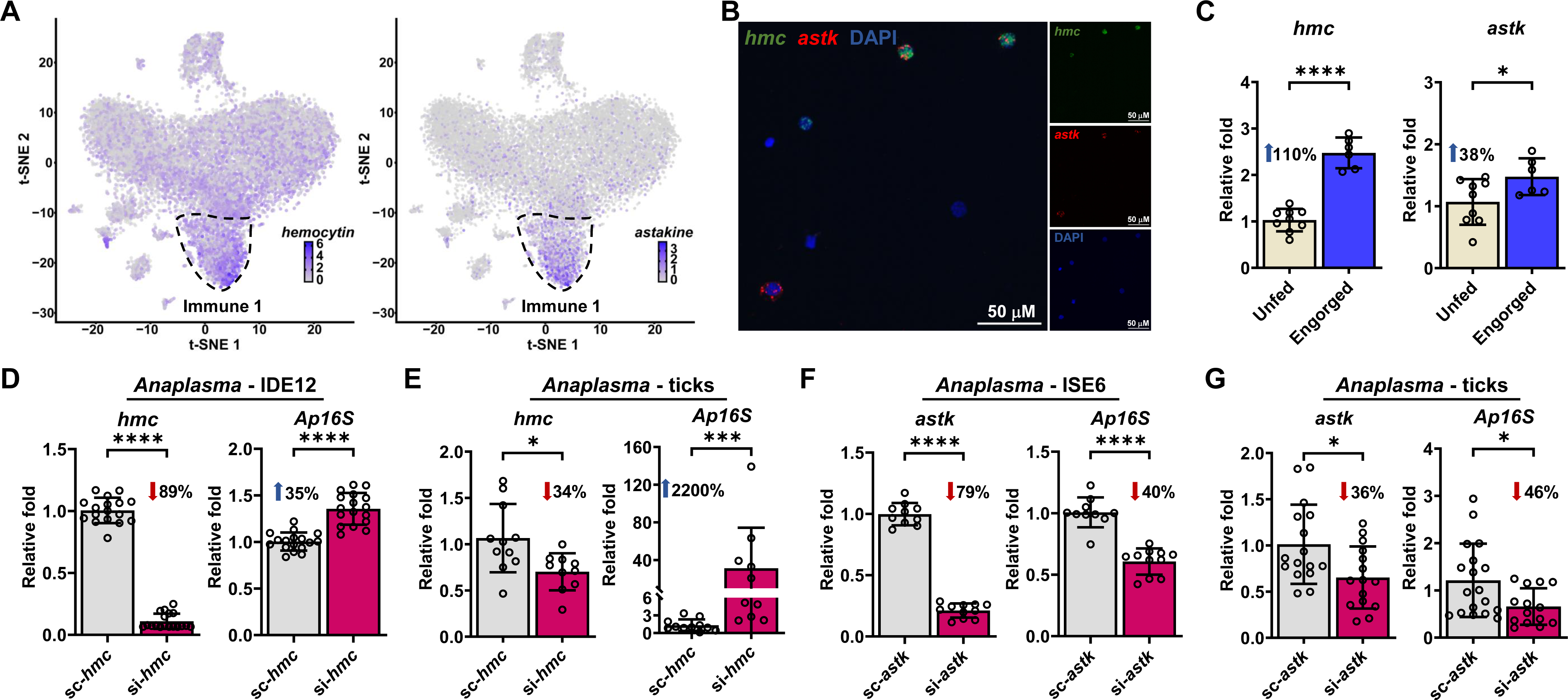
Hemocytin (hmc) and astakine (astk) affect A. phagocytophilum infection. **(A)** Expression of *hmc* (left) and *astk* (right) on t-Distributed Stochastic Neighbor Embedding (t-SNE) plots of hemocytes collected from engorged nymphs, with their highest expression denoted in the Immune 1 cluster (outlined). **(B)** RNA FISH of *I. scapularis* hemocytes probed for *hmc* (green), *astk* (red), and nuclei (DAPI). **(C)** Expression of *hmc* (left) and *astk* (right) in hemocytes from unfed (ivory) or engorged (dark blue) ticks were evaluated by RT-qPCR (*n*=6-9; samples represent 40-80 pooled ticks). **(D)** *hmc* (left) silencing efficiency and *A. phagocytophilum* burden (right) in IDE12 cells. Cells were transfected with *hmc* siRNA (si-*hmc*) or scrambled RNA (sc-*hmc*) for seven days prior to *A. phagocytophilum* infection (*n*=17-18). **(E)** *hmc* silencing efficiency (left) and bacterial acquisition (right) in ticks microinjected with si-*hmc* or sc-*hmc* fed on *A. phagocytophilum*-infected mice (*n*=10-11). **(F)** *astk* silencing efficiency (left) and *A. phagocytophilum* burden (right) in ISE6 cells. Cells were transfected with *astk* siRNA (si- *astk*) or scrambled RNA (sc-*astk*) for seven days prior to *A. phagocytophilum* infection (*n*=10- 11). **(G)** *astk* silencing efficiency (left) and bacterial acquisition (right) in ticks microinjected with si-*astk* or sc-*astk* fed on *A. phagocytophilum*-infected mice (*n*=14-18). Bacterial burden was quantified by *A. phagocytophilum 16srRNA* (*Ap16S*) expression. Results are represented as mean ± SD. At least two biological replicates were performed. Statistical significance was evaluated by an **(C-D, F-G)** unpaired t-test with Welch’s correction or **(E)** Mann–Whitney U test. \**p*<0.05; \*\*\**p*<0.001; \*\*\*\**p*<0.0001.

Bulk RNA-seq of hemocytes originating from the tick *Amblyomma maculatum* showed differential expression of *hemocytin* and *astakine* during infection with the intracellular bacterium *Rickettsia parkeri* (*31*). Our scRNA-seq revealed downregulation of *hemocytin* and *astakine* expression during *A. phagocytophilum* and *B. burgdorferi* infection (Figure 3D, Supplementary Dataset 9). We continued investigating the role of *hemocytin* and *astakine* during *A. phagocytophilum* infection of *I. scapularis*. We utilized small-interfering RNA (siRNA) to knockdown gene expression in tick cells and *I. scapularis* nymphs, as genome editing through CRISPR has only been introduced for morphological assessment in ticks (*32*). Accordingly, tick cells and *I. scapularis* nymphs were treated with siRNAs targeting *hemocytin* or *astakine* before infection with *A. phagocytophilum*. While we observed a significant increase in bacterial load in *hemocytin*-silenced treatments (Figure 4D-E), a decrease in infection was measured after *astakine* silencing (Figure 4F-G). Importantly, no differences in attachment were observed between conditions (Supplementary Figure 9). Overall, we observed contrasting effects of *hemocytin* or *astakine* on *A. phagocytophilum* infection, highlighting the complexity of the tick immune system.

### *Hemocytin* affects the JNK pathway of *I. scapularis*

Arachnids constitutively express AMPs in hemocytes (*7, 33, 34*). However, how immune signaling cascades are regulated in these species, including ticks, remains obscure. Given that we detected higher *A. phagocytophilum* load when we silenced *hemocytin* in ticks (Figure 4D-E), we asked whether hemocytin affects immune signaling pathways in *I. scapularis*. We focused on the IMD and the c-Jun N-terminal kinase (JNK) pathway, as previous reports indicated the involvement of these molecular networks during bacterial exposure (*6, 7*).

After transfection with siRNA targeting *hemocytin,* we found a decrease in JNK phosphorylation in *hemocytin*-silenced tick cells, without alteration in Relish cleavage (Figure 5A-B). To complement our findings, we overexpressed *hemocytin* in tick cells through CRISPR activation (CRISPRa). CRISPRa has been widely used to enhance the expression of an endogenous locus, employing a catalytically inactive Cas9 (dCas9) fused with transcription activators and single guide RNAs (sgRNAs) that direct the modified enzyme to the promoter region of a gene of interest (*35*). However, so far none of the CRISPRa effectors has been tested in tick cell lines and no endogenous promoter has been identified for genetic expression of sgRNAs. We optimized reagents and developed a protocol to up-regulate *hemocytin* in ISE6 cells using two rounds of nucleofection with different expression plasmids: one expressing dCas9-VPR paired with neomycin resistance under the control of the CMV promoter and the second expressing either a sgRNA specific to the promoter region of *hemocytin* (*hmc*-sgRNA) or a scrambled sgRNA (sc-sgRNA) driven by an endogenous RNA polymerase III promoter (Figure 5C). Tick ISE6 cells were used as a platform for CRISPRa given the lower expression of *hemocytin* compared to IDE12 cells (Supplementary Figure 10). Strikingly, we detected a 277% increase in the expression of *hemocytin* in dCas9^+^ cells transfected with the *hmc*-sgRNA compared to dCas9^+^ cells transfected with the sc-sgRNA (Figure 5D). Importantly, we also noticed elevated levels of *jun* (the transcription factor for the JNK pathway) and *jnk* expression in the *hmc*-sgRNA treatment (Figure 5D).

**Figure 5:**
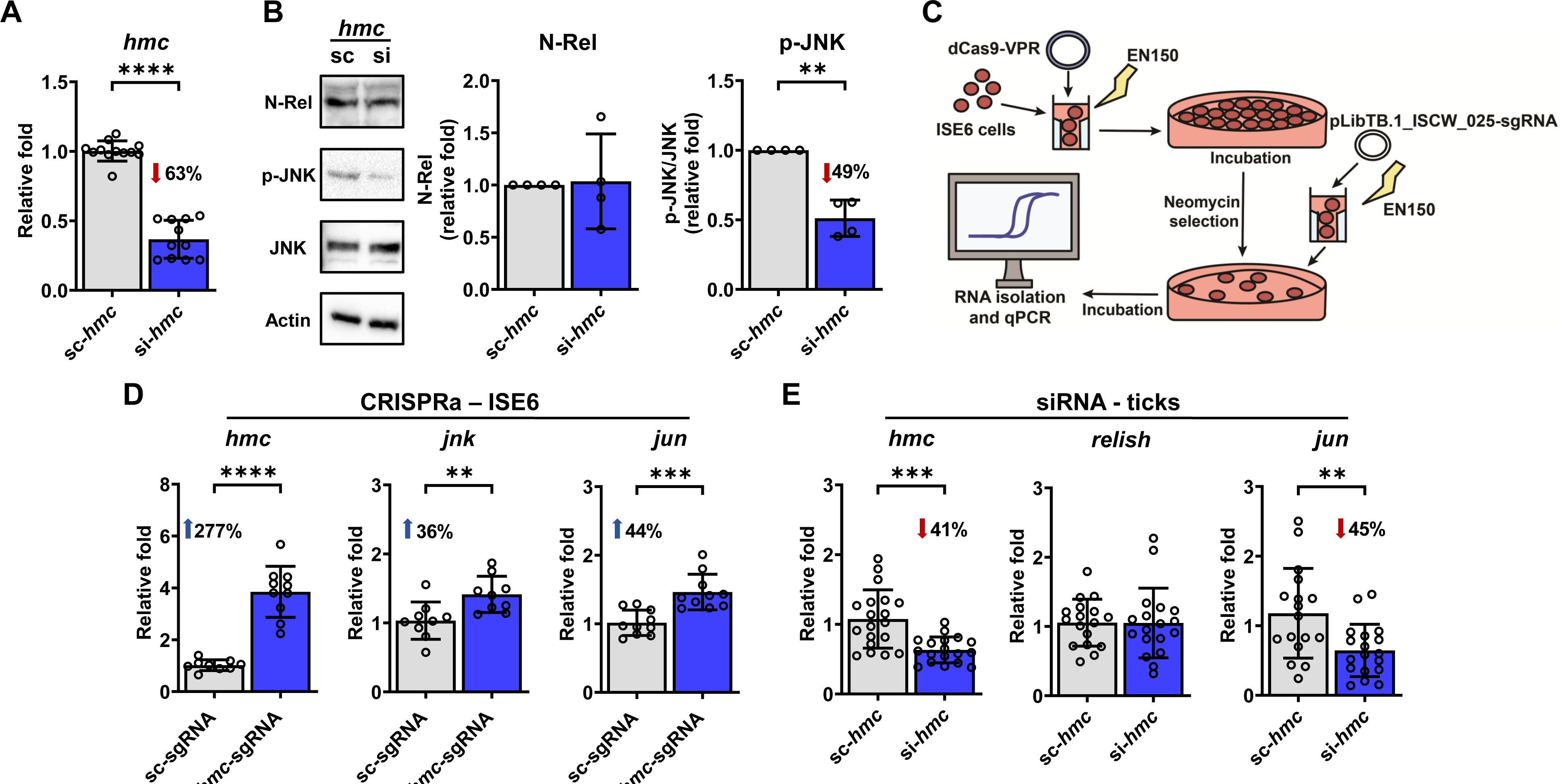
*Hemocytin* (*hmc*) positively impacts the JNK pathway in *I. scapularis*. (A) Cells were transfected with *hmc* siRNA (si-*hmc*) or scrambled RNA (sc-*hmc*) (*n*=11-12). *hmc* silencing efficiency in IDE12 cells. **(B)** Representative western blot (left) of N-Rel and p- JNK during treatment with sc-*hmc* (lane 1) or si-*hmc* (lane 2). N-Rel and p-JNK protein expression was quantified (right) in si-*hmc* (blue) or sc-*hmc* (grey) IDE12 cells. For data normalization, values were divided by the scrambled control value. N-Rel values are normalized to Actin and p-JNK values are normalized to JNK. Western blot images show one representative experiment out of four. **(C)** Schematic of CRISPRa overexpression of *hmc* in ISE6 cells. **(D)** Expression of *hmc* (left) *jnk* (middle) and *jun* (right) in dCas9^+^ISE6 cells transfected with the *hmc*-sgRNA (blue) compared with the scrambled-sgRNA (grey) evaluated by RT-qPCR (*n*=9- 10). **(E)** *hmc* (left), *relish* (middle) and *jun* (right) expression in ticks microinjected with *hmc* siRNA (si-*hmc*; blue) or scrambled RNA (sc-*hmc*; grey) fed on uninfected mice (*n*=17- 19). Results are represented as mean ± SD. At least two biological replicates were performed. Statistical significance was evaluated by an unpaired t-test with Welch’s correction. \*\**p*<0.01; \*\*\**p*<0.001; \*\*\*\**p*<0.0001. N-Rel = cleaved Relish; p-JNK = phosphorylated JNK; JNK = c-Jun N-terminal kinase.

To corroborate our results *in vivo*, we microinjected unfed ticks with a scrambled control or siRNA targeting *hemocytin* and allowed them to feed on naïve mice. Upon repletion, we measured the expression of *jun* and *relish*. Consistently, we found that reduction in *hemocytin* expression led to a decrease in *jun* levels in ticks, without affecting *relish* expression (Figure 5E). Notably, the alteration in the JNK signaling pathway by hemocytin in ticks was unrelated to hemocyte proliferation, differentiation or phagocytosis (Supplementary Figures 11-12). Overall, we uncovered a role for hemocytin in the activation of the JNK pathway in *I. scapularis*.

### Astakine induces hemocyte proliferation and differentiation in *I. scapularis*

We found that hematophagy induces hemocyte proliferation and differentiation (Figure 1C) while upregulating *astakine* expression (Figure 4C). Thus, we investigated whether *astakine* was directly implicated in the proliferation or differentiation of *I. scapularis* hemocytes. We microinjected increasing amounts of recombinant astakine (rAstk) in unfed nymphs and observed a dose-dependent increase in the total number of hemocytes (Figure 6A). Specifically, we measured a decrease in the percentage of prohemocytes and an increase in plasmatocytes (Figure 6B). These findings were in accordance with alterations observed during normal blood-feeding (Figure 1C). Interestingly, we also detected an increase in tick IDE12 cell numbers *in vitro* following treatment with rAstk, supporting a role of astakine in inducing cell proliferation in hemocyte-like cells (Supplementary Figure 13). To support these results, we then silenced *astakine* by microinjecting unfed nymphs with siRNA before feeding on uninfected mice. We recovered 41% less hemocytes and observed an increase in the percentage of prohemocytes with a decrease in plasmatocytes from engorged ticks when *astakine* was silenced (Figure 6C- E). Collectively, we determined that astakine acts on hematopoietic processes in the ectoparasite *I. scapularis*.

**Figure 6:**
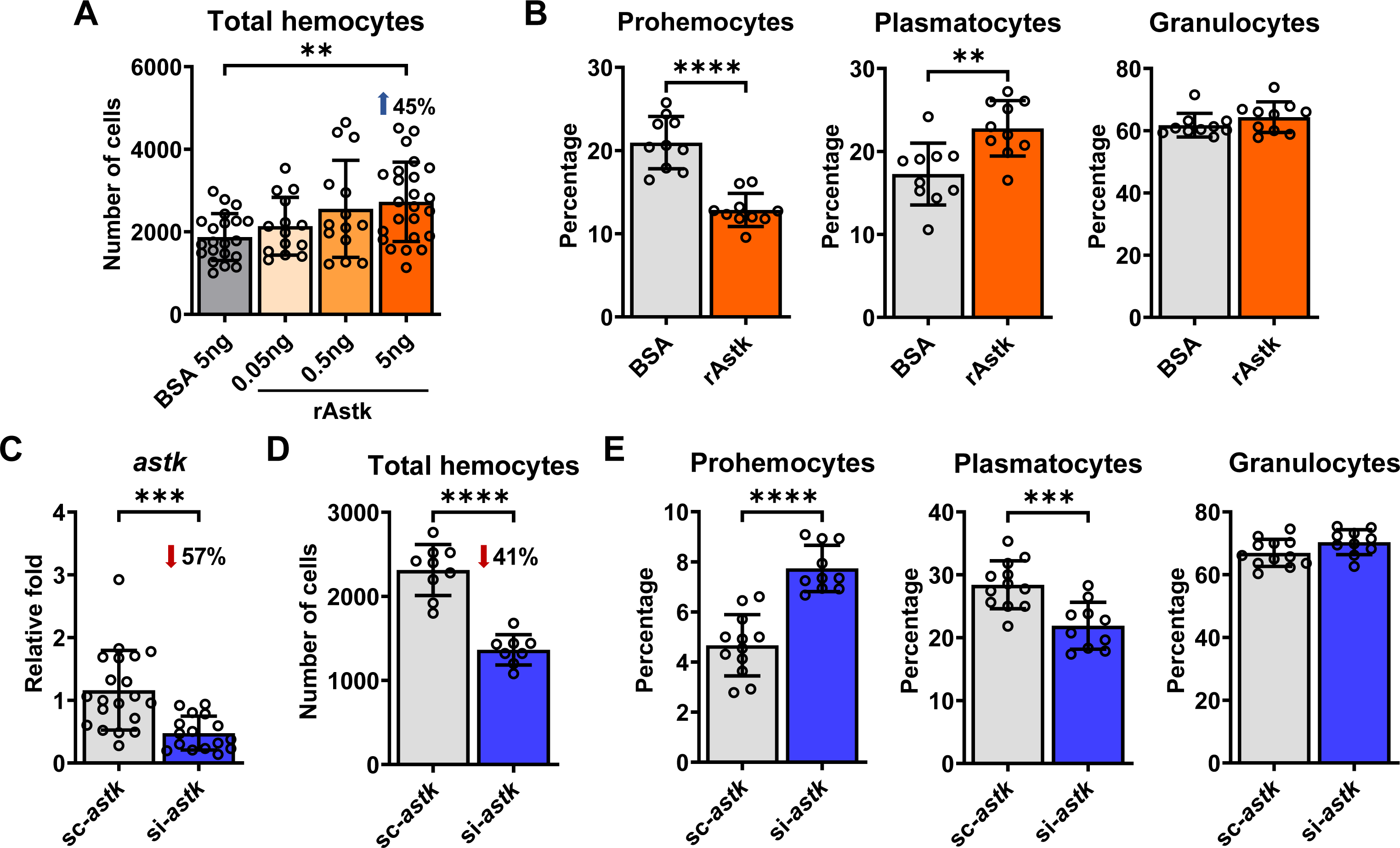
Astakine (astk) induces hemocyte proliferation and differentiation in *I. scapularis*. **(A)** Total number of hemocytes collected from unfed *I. scapularis* nymphs microinjected with corresponding amounts of rAstk (orange) or bovine serum albumin (BSA; grey) as a control (*n*=14-24). **(B)** Percentage of hemocyte morphotypes present in unfed *I. scapularis* nymphs microinjected with 5ng rAstk (orange) compared to BSA controls (grey) (*n*=10). **(C)** *astk* silencing efficiency, **(D)** total number of hemocytes and **(E)** percentage of hemocyte morphotypes in engorged ticks previously microinjected with *astk* siRNA (si-*astk*; blue) or scrambled RNA (sc-*astk*; grey) and fed on uninfected mice (*n*=8-20). Results represent mean ± SD. At least two biological replicates were performed. Statistical significance was evaluated by **(A)** one-way ANOVA or **(B-E)** an unpaired t-test with Welch’s correction. \*\**p*<0.01; \*\*\**p*<0.001; \*\*\*\**p*<0.0001. rAstk=recombinant astakine.

### Manipulation of hemocyte subtypes or marker genes affects tick physiology

The non- canonical functions of hemocytes in arthropods are underappreciated. We observed an increased expression of genes and expansion of hemocyte subtypes involved in metabolism, cell proliferation and development following a blood meal (Figures 1-2), suggesting participation of these immune cells in the feeding or molting process. To investigate this observation, we manipulated hemocyte numbers in *I. scapularis* using clodronate liposomes (CLD), which have been used to deplete phagocytic hemocytes in flies, mosquitoes and, more recently, ticks (*31, 36, 37*). We found that the CLD treatment reduced the total number of hemocytes in nymphs (Figure 7A). We detected an increase in the proportion of granulocytes together with a decrease in the percentages of prohemocytes and plasmatocytes compared to ticks treated with empty liposomes (Figure 7B). These results indicated that CLD affects phagocytic hemocytes in *I. scapularis*.

**Figure 7:**
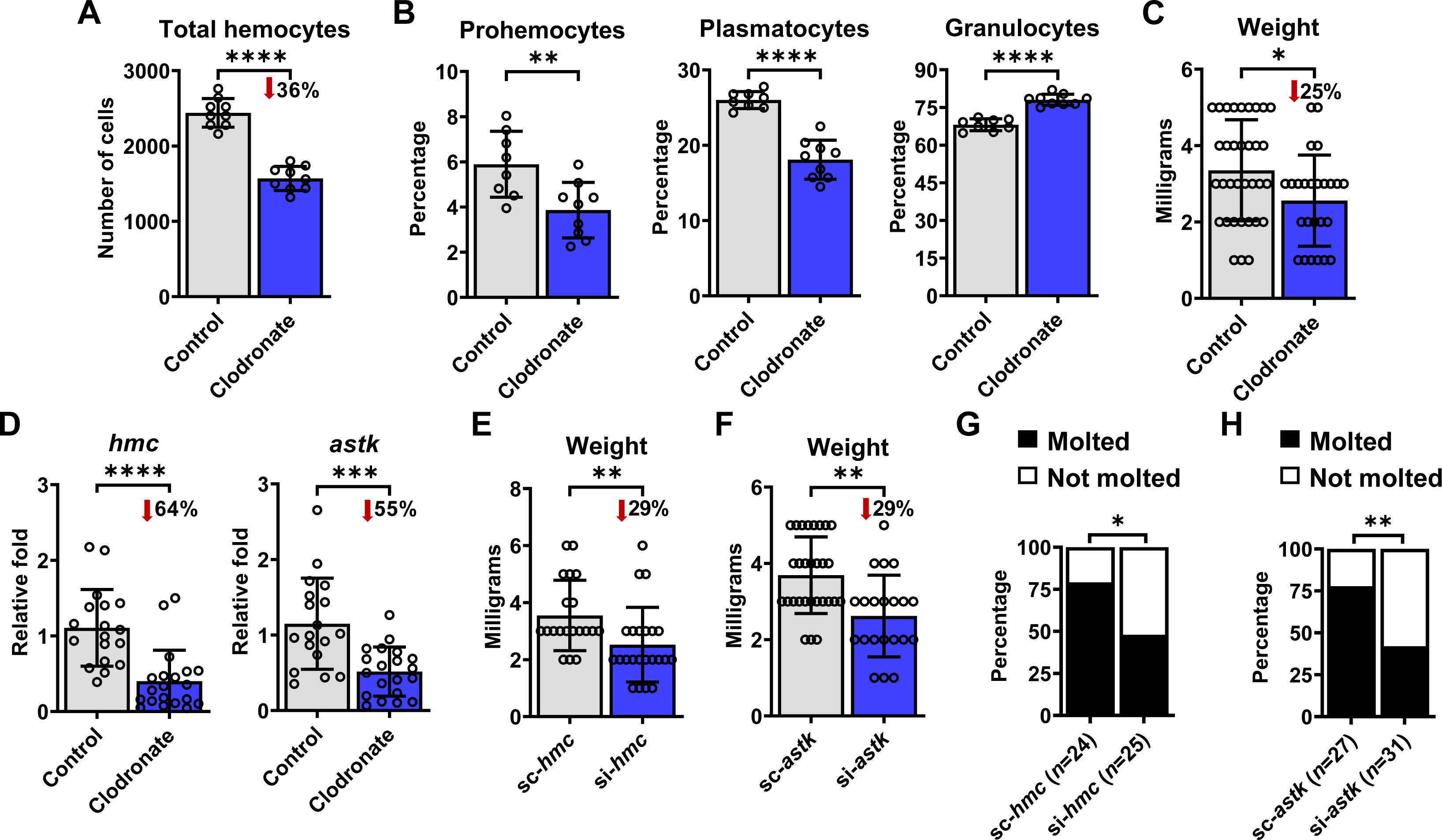
Manipulation of hemocyte subtypes and their marker genes affects tick fitness. **(A-D)** Ticks were microinjected with clodronate (blue) or empty liposomes as a control (grey) and allowed to feed on uninfected mice. **(A)** Total number of hemocytes (*n*=9) and **(B)** morphotype percentages were determined in the hemolymph (*n*=8-9). **(C)** Weight of engorged nymphs (*n*=25-34) and **(D)** expression of *hemocytin* (*hmc*) and *astakine* (*astk*) in individual ticks was measured (*n*=18-20). **(E)** Weight of engorged nymphs post-microinjection with *hmc* siRNA (si-*hmc*; blue) or scrambled RNA (sc-*hmc*; grey) fed on uninfected mice (*n*=20-23). **(F)** Weight of engorged nymphs microinjected with *astk* siRNA (si-*astk*; blue) or scrambled RNA (sc-*astk*; grey) fed on uninfected mice (*n*=21-29). **(G)** Percentage of nymphs that molted to adults after treated with si-*hmc* or sc-*hmc* and **(H)** si-*astk* or sc-*astk*. Results are represented as **(A-F)** mean ± SD or as **(G-H)** a percentage of ticks that molted from the total recovered after feeding. At least two biological replicates were performed. Statistical significance was evaluated by **(A-B, D)** an unpaired t-test with Welch’s correction, **(C, E, F)** Mann–Whitney U test or **(G-H)** by a Fisher exact test. \**p*<0.05; \*\**p*<0.01; \*\*\**p*<0.001; \*\*\*\**p*<0.0001.

To determine the contribution of tick hemocytes to the feeding process, we then used CLD to manipulate immune cell populations. Although no differences in attachment were observed (Supplementary Figure 14), engorged ticks treated with CLD weighed significantly less after feeding, suggesting that phagocytic hemocytes are vital for proper tick hematophagy (Figure 7C). Notably, expression of *hemocytin* and *astakine* was also decreased after CLD treatment, implying that the Immune 1 cluster is likely phagocytic (Figure 7D). Thus, we postulated that altering the expression of Immune 1 marker genes should impact tick hematophagy.

Accordingly, we observed a significant decrease in weight in ticks silenced for *hemocytin* or *astakine* fed on either uninfected or *A. phagocytophilum*-infected mice, without differences in tick attachment (Figure 7E-F; Supplementary Figure 15).

Given the close relationship between blood feeding and ecdysis in ticks, we next posited that *hemocytin* and/or *astakine* might be involved in molting of *I. scapularis* nymphs. To test this hypothesis, we microinjected nymphs with scRNA or siRNA targeting *hemocytin* or *astakine* and allowed them to engorge on naïve mice, and subsequently molt into adults. Molting success was significantly lower in both gene-silenced groups as compared to scrambled controls (Figure 7G). Overall, the data indicate that tick hemocyte subtypes and associated marker genes pleiotropically impact tick immunity, feeding and molting.

## Discussion

The immune response against microbial infection has been extensively studied in Dipteran insects, but these species do not fully account for the variation in fundamental processes that exists in arthropods (*4–10, 18*). In this study, we expand on the previous morphological classification of tick hemocytes. We characterized hemocytes at single-cell resolution in combination with orthogonal approaches to uncover hemocyte subtypes and molecular markers in *I. scapularis*. The data revealed immune and metabolic alterations of tick hemocytes in response to blood feeding and microbial infection. In contrast to mosquitoes (*38–40*), for *I. scapularis* ticks, we noticed the emergence of new hemocyte clusters and distinct biological signatures after hematophagy.

Hemocytes with immune-related roles exhibited overrepresentation of genes involved in peptide secretion, agglutination and clotting, extracellular matrix remodeling and protein degradation. Thus, it is conceivable that these clusters not only participate in antimicrobial responses, but also aid in the extensive internal remodeling of tissues needed during blood meal acquisition (*22*). We also uncovered hemocytes specialized towards cellular proliferation. These hemocytes expressed genes involved in ecdysteroid biosynthesis, which is essential for regulating molting and development (*7, 41*). Metabolic hemocytes were enriched in solute transport, lipid and protein metabolism, histamine binding and detoxification genes. These molecular features suggested an involvement in the feeding process by metabolizing nutrients and xenobiotics present in the blood meal.

Ticks have co-evolved a non-lethal relationship with *A. phagocytophilum* and *B. burgdorferi*.

Neither microbe typically results in morbidity or mortality in *I. scapularis* ticks (*2, 3*). Our work presents an opportunity for comparative biology in arthropods as to why tick infection with *A. phagocytophilum* or *B. burgdorferi* does not induce expansion of new hemocyte subtypes.

Avoidance of strong cellular responses highlights the discrete balance between immunity and survival that has been evolutionarily conserved among hematophagous arthropods.

We established molecular programs in hemocytes associated with bacterial infection of *I. scapularis*. We mechanistically characterized two marker genes: *hemocytin* and *astakine*.

*Drosophila* mutants for *hemolectin,* the ortholog of *hemocytin* in flies, display only mild phenotypic defects (*24, 25*). In contrast, silencing of *hemocytin* in ticks led to an elevated *A. phagocytophilum* infection during hematophagy independent of phagocytosis. Additionally, we uncovered that *hemocytin* positively affects the tick JNK pathway. Thus, we posit that *hemocytin* balances tick homeostatic processes during microbial infection and physiological alterations.

Similar to what has been observed in crustaceans (*27*), we uncovered a function for *astakine* in hematopoiesis of ticks. Prior evidence suggested that blocking phagocytosis in hemocytes reduces *A. phagocytophilum* dissemination to other tick tissues (*42*). We found that silencing *astakine* resulted in a decreased *A. phagocytophilum* load in *I. scapularis,* which correlated with a diminished number of phagocytic hemocytes. Future studies determining the cellular source, receptors and signaling pathways of astakine will reveal mechanisms driving immune proliferation and differentiation in ticks.

Hemocytes are important for organogenesis and development, clearing of apoptotic cells and tissue communication (*43, 44*). We showed that alterations in hemocyte subtypes and *astakine* and *hemocytin* levels affected fitness in *I. scapularis*. Given the transcriptional profiles induced by feeding and the circulatory nature of hemocytes, we hypothesize that these cells are involved in the internal reorganization of tissues and organs needed during hematophagy. In contrast to *astakine* knockdown or the CLD treatment, *hemocytin* knockdown disrupted normal tick feeding without altering hemocyte numbers or their phagocytic capacity. As silencing *hemocytin* alters the JNK signaling network, we posit that the JNK pathway in ticks is implicated in organismal growth. We observed defects in tick molting when hemocyte subtypes and *astakine* and *hemocytin* levels were altered. As opposed to insects, Acari rely solely on the synthesis of ecdysteroids derived from cholesterol for molting (*7, 41*). Our results suggest the involvement of hemocytes in ecdysteroid synthesis and transport as homologs of most genes related to lipid metabolism were altered after a blood meal. Studying strict hematophagy in ticks enables the discovery of non-canonical roles of immune cells shaped by adaptations to prolonged feeding.

In summary, we leveraged the power of systems biology in combination with reductionist approaches to shed light on the complexity and dynamic attributes of biological processes associated with immune cells of an obligate hematophagous ectoparasite. Our findings uncovered canonical and non-canonical roles of tick hemocytes, which might have evolved as a consequence of evolutionary associations of *I. scapularis* with its vertebrate hosts and microbes they interact. In the future, small- and large-scale approaches described in this study could be adapted to serve as a foundation to construct an integrated atlas of the medically relevant tick *I. scapularis*.

## Materials and Methods

### Reagents and resources

All primers, reagents, resources, and software used in this study, together with their manufacturers information and catalog numbers are listed in Supplementary Tables 1 and 2.

### Cell lines

The *I. scapularis* IDE12 and ISE6 cell lines were obtained from Dr. Ulrike Munderloh at University of Minnesota. Cells were cultured in L15C300 medium supplemented with 10% heat inactivated fetal bovine serum (FBS, Sigma), 10% tryptose phosphate broth (TPB, Difco), 0.1% bovine cholesterol lipoprotein concentrate (LPPC, MP Biomedicals) at 34°C in a non-CO2 incubator. ISE6 cells were grown to confluency in capped T25 flasks (Greiner bio-one) and either seeded at 3x10^6^ cells/well in 6-well plates (Millipore Sigma) or 3x10^5^ cells/well in 24-well plates (Corning). IDE12 cells were also grown to confluency in T25 flasks and either seeded at 1x10^6^ cells/well in 6-well plates (Millipore Sigma) or 3x10^5^ cells/well in 24-well plates (Corning). The human leukemia cell line, HL-60, was obtained from ATCC and maintained in RPMI-1640 medium with L-Glutamine (Quality Biological) supplemented with 10% FBS (Gemini Bio- Products) and 1% GlutaMax (Gibco), in vented T25 flasks (CytoOne) at 37°C and 5% CO2. All cell cultures were confirmed to be *Mycoplasma*-free (Southern Biotech).

### Bacteria, ticks, and mice

*A. phagocytophilum* strain HZ was grown as previously described in HL-60 cells at 37°C, using RPMI medium supplemented with 10% FBS and 1% Glutamax (*45*). Bacterial numbers were calculated using the following formula: number of infected HL-60 cells × 5 morulae/cell × 19 bacteria/morulae × 0.5 recovery rate (*46*). To determine the percentage of infection, cells were spun onto microscope slides using a Epredia™ Cytospin™ 4 Cytocentrifuge (Thermo Scientific), stained using the Richard-Allan Scientific™ three-step staining (Thermo Scientific) and visualized by light microscopy using a Primo Star™ HAL/LED Microscope (Carl Zeiss). For *in vitro* experiments, bacteria were purified in L15C300 media by passing infected cells through a 27-gauge bent needle and centrifugation, as previously described (*45*). Isolated bacteria were inoculated onto tick cells at a multiplicity of infection (MOI) of 50 for 48 hours. For *in vivo* experiments, a total of 1x10^7^ *A. phagocytophilum*-infected cells/mL was resuspended in 100 µL of 1X phosphate-buffered saline (PBS) and intraperitonially injected into mice. Infection progressed for 6 days prior to tick placement.

*B. burgdorferi* B31 clone MSK5 (passage 3-5) was cultured in Barbour-Stoenner Kelly (BSK)-II medium supplemented with 6% normal rabbit serum (Pel-Freez) at 37°C. Before mouse infection, PCR plasmid profiling was performed as described elsewhere (*47*). To determine infectious inoculum, spirochetes were enumerated under a 40X objective lens using a dark-field microscope (Carl Zeiss™ Primo Star™ Microscope) by multiplying the number of bacteria detected per field x dilution factor x 10^6^ (*45, 47*). For intradermal inoculation, spirochetes were washed, resuspended in 50% normal rabbit serum at a concentration of 1x10^6^ *C. burgdorferi/*mL and anesthetized C3H/HeJ mice were injected with 100 µL of inoculum (1x10^5^ total *B. burgdorferi*). Infected mice were maintained for at least 14 days prior to tick placement.

*I. scapularis* nymphs were obtained from Oklahoma State University and University of Minnesota. Upon arrival, ticks were maintained in a Percival I-30BLL incubator at 23°C with 85% relative humidity and a 12/10-hours light/dark photoperiod regimen. Age matched (6-10 weeks), male C57BL6 and C3H/HeJ mice were supplied by Jackson Laboratories or the University of Maryland Veterinary Resources. For all tick experiments, capsules were attached to the back of mice using a warmed adhesive solution made from 3 parts gum rosin (Millipore- Sigma) and 1 part beeswax (Thermo Scientific) 24 hours prior to tick placement. All experiments were done using C57BL6 mice, except for the ones including *B. burgdorferi* infection, in which the mice genetic background was homogenized to the C3H/HeJ. Mouse breeding, weaning and experiments were performed under guidelines from the NIH (Office of Laboratory Animal Welfare (OLAW) assurance numbers A3200-01) and pre-approved by the Institutional Biosafety (IBC-00002247) and Animal Care and Use (IACUC-0119012) committee of the University of Maryland School of Medicine.

### Hemocyte collection

Hemolymph was collected from *I. scapularis* nymphs in L15C300 + 50% FBS on ice from individual unfed, partially fed or engorged ticks placed on uninfected, *A. phagocytophilum-* or *B. burgdorferi*-infected mice, using non-stick RNAase-free 1.5 mL microtubes (Ambion) and siliconized pipet tips. Briefly, ticks were immobilized on a glass slide using double-sided tape (3M) and covered in a sphere of 10 µL of L15C300 + 50% FBS. Forelegs were incised at the tarsus, and forceps were used to gently apply pressure to the tick body to release hemolymph into the media. Enumeration of total hemocytes was determined for each tick using a Neubauer chamber. For differentiation of hemocyte morphotypes, 100 µL of L15C300 was added in a microtube for each tick before preparation on Fisherbrand™ Superfrost™ Plus microscope slides (Fisher Scientific) using a Epredia™ Cytospin™ 4 Cytocentrifuge (Thermo Scientific).

Afterwards, hemocytes were stained using the Richard-Allan Scientific™ three-step staining (Thermo Scientific) and evaluated morphologically under a Primo Star™ HAL/LED Microscope (Carl Zeiss). Images were acquired using an Axiocam 305 color camera (Carl Zeiss) with the ZEN software (Carl Zeiss).

For RNA extraction, hemocytes were collected as mentioned above using PBS (4 µL for unfed and 6 µL for engorged ticks) instead of media. For each biological replicate, hemocytes from 80 unfed or 40 engorged ticks were pooled in 700 µL of TRIzol.

### Bulk RNA sequencing

Three independent hemocyte collections were performed for each condition (unfed and engorged), as mentioned above. After collection, RNA was extracted from TRIzol following the manufacturer’s instruction with the addition of 1 µg of glycogen per sample. The RNA integrity (RIN) was assessed for each sample using a Eukaryote Total RNA Nano Chip Assay on a Bioanalyzer (Agilent). RIN values ranged between 7.6 to 9.4. Strand-specific, dual unique indexed libraries for sequencing on Illumina platforms were generated using the NEBNext® Ultra™ II Directional RNA Library Prep Kit for Illumina® (New England Biolabs). The manufacturer’s protocol was modified by diluting adapters 1:30 and using 3 µL of this dilution. The size selection of the library was performed with SPRI-select beads (Beckman Coulter Genomics). Glycosylase digestion of the adapter and 2^nd^ strand was performed in the same reaction as the final amplification. Sample input was polyA-enriched. Libraries were assessed for concentration and fragment size using the DNA High Sensitivity Assay on the LabChip GX Touch (Perkin Elmer). The resulting libraries were sequenced on a multiplexed paired end 100bp Illumina NovaSeq 6000 S4 flowcell, generating an average of 92 million read pairs per sample. RIN analysis, library preparation and sequencing were completed by Maryland Genomics at the Institute for Genome Sciences, University of Maryland School of Medicine.

One sample was excluded from subsequent analysis due to considerably smaller library size compared to the rest.

After sequencing, all reads were aligned to the *I. scapularis* genome (GCF_016920785.2_ASM1692078v2) using HISAT v2.0.4 (*18, 48*). Next, we potential PCR duplicates were removed by keeping only one of the read pairs mapped to the same genomic location of the same strand and then tallied the number of “unique reads” mapped to each annotated *I. scapularis* gene. The generated count matrix was used as input for differential expression analysis with edgeR v3.36.0 (*49*). During data normalization, only transcripts with at least 10 counts per million (cpm) in two or more samples were kept for downstream analysis. Differentially expressed genes (DEGs) were defined based on the following thresholds: (1) a false discovery rate (FDR) <0.05, and (2) -0.5146<LOGFC>0.3785, corresponding to at least a 30% reduction or increase in gene expression. All sequencing reads were deposited into the NCBI Sequence Read Archive under the BioProject accession PRJNA906572.

### Single-cell RNA sequencing

For single-cell RNA sequencing (scRNA-seq), hemocytes were collected from individual unfed, engorged, or *A. phagocytophilum-* or *B. burgdorferi-*infected ticks and pooled in 300 µL of L15C300 + 50% FBS (*n*= 90 for unfed and *n*=50 for engorged ticks). Cells were filtered through a 40 µm cell strainer (Millipore Sigma) and enriched for live cells using OptiPrep density gradient centrifugation (*50*). Briefly, 200-300 µL of the filtered solution was overlaid onto 2 mL of OptiPrep™ Density Gradient Medium (1.09 g/mL in 1X PBS; Sigma) and centrifugated at 1,000xg for 10 minutes at 4°C, using a swinging bucket without breaking. After centrifugation, 200 µL of the interphase was collected in a new microtube and spun at 1,000xg for 10 min at 4°C. Cells were concentrated in 30 µL of 1X PBS + 0.2 U RNasin® Plus Ribonuclease Inhibitor (Promega), from which 5 µL were used for measuring viability and cell numbers by trypan blue exclusion assay. The final concentration of cells ranged from 350 to 500 cells/µL, with viability ranging between 95.7% and 98.5%.

An estimated 5,000-7,000 cells per condition were loaded on a 10X Genomics Chromium controller and four individually barcoded 3’-end scRNA-seq libraries were prepared according to the manufacturer’s protocol. Each library was then sequenced on an Illumina high-output sequencer to generate a total of 816,376,790 75-bp paired-end reads. To process all scRNA-seq reads, a custom pipeline previously used to analyze single-cell data from *Plasmodium* parasites similar to 10X Genomics Cell Ranger software was adapted (*51*). First, only reads longer than 40 bp after trimming any sequences downstream of 3’ polyadenylation tails were kept. These reads were then mapped to the *I. scapularis* genome (GCF_016920785.2_ASM1692078v2) using HISAT version 2.0.4 (*18, 48*). Out of the mapped reads, “unique reads” were identified by keeping only one of the reads that had identical 16-mer 10X Genomics barcode, 12-mer unique molecular identifier, and mapped to the same genomic location of the same strand. Finally, the 10X Genomics barcodes were used to separate reads derived from each individual cell and count the number of unique reads mapped to each annotated *I. scapularis* gene (from transcription start site to 500 bp after the annotated 3’-end). All sequencing reads are deposited into the NCBI Sequence Read Archive under the BioProject accession PRJNA905678 and the code for mapping is available through the GitHub website (https://github.com/Haikelnb/scRNASeq_Analysis) The scRNA-seq libraries were combined into a single dataset using scran v1.26.2 (*52*).

Genes containing “40s_ribosomal” and “60s_ribosomal” were removed from the entire dataset. Based on Principal Component Analysis (PCA), unfed and fed hemocyte samples were analyzed independently for downstream analyses. Low quality cells were removed with the following thresholds: less than 600 unique reads, less than 150 mapped genes, and 30% or higher mitochondrial transcripts. The remaining transcriptomes were normalized by first calculating size factors via scran functions quickCluster and computeSumFactors and computing normalized counts for each cell with logNormCounts function in scater v1.22.0 (*53*). For downstream analysis, highly variable genes were selected using getTopHVGs before performing PCA and t-distributed Stochastic Neighbor Embedding (t-SNE) projections.

Clustering was conducted using a kmeans value of 20. Differential gene expression between clusters was calculated using the find marker function in scran v1.26.2 (*52*). The R package slingshot v2.2.1 was used to perform pseudotime inference where trajectories began from the “Proliferative 1” cluster, based on predicted function (*54*). To determine differences in gene expression between each infected condition compared to the reference factor (uninfected), clusters were grouped based on annotated function (e.g. metabolism, proliferation, immune) and MAST v1.24.1 was used to test for significance under the Hurdle model adjusting for the cellular detection rate (*55*). The code for the analysis is available through the GitHub website (https://github.com/HLaukaitisJ/PedraLab_hemocyte_scRNAseq).

### Functional annotation and enrichment analysis

Functional annotation and Gene Ontology Enrichment analysis were done through VectorBase, using the LOC number of all DEGs (bulk RNA-seq) or marker genes (scRNA-seq) as input, and the following parameters: (1) Organism: “*Ixodes scapularis* PalLabHiFi”; (2) Ontology: “Molecular Function” or “Biological Process”; (3) adjusted p-value cutoff (Benjamini) < 0.05 (*56*). Additionally, the putative function of the top 50 marker genes in each single cell cluster were manually assigned based on 10 categories (“ncRNA/Pseudogene”, “detoxification”, “secreted protein/extracellular matrix”, “metabolism”, “immunity”, “hormone related”, “cell proliferation/differentiation”, “protein synthesis”, “actin polymerization/cell rearrangement”, “unknown”). First, the protein coding sequence of each gene was retrieved from the NCBI website. Next, homologues from each gene were searched in the *D. melanogaster* genome using the BLAST tool from Flybase (*57*). If a match was found with an E value < 10^-20^, then the genes were assumed to have some (E value was between < 10^-20^ to 10^-39^) or complete (E value < 10^-40^) homology. The function of the *I. scapularis* gene was interpreted based on the description for *D. melanogaster*. Comparison to gene expression profiles in specific *D. melanogaster* hemocyte subtypes was assessed using the DRSC/TRiP Functional Genomics Resources dataset (*50*). Alternatively, if a high match to known *D. melanogaster* genes was not found, then the complete coding sequence of the *I. scapularis* gene was used to search for protein domains using InterPro and the functional information retrieved from Pubmed or UniProt (*58, 59*). Otherwise, the gene function was assigned as “unknown”.

### RNA-fluorescence *in situ* hybridization (FISH)

The expression of specific genes in hemocytes was determined by RNA-FISH using the RNAScope-Multiplex Fluorescent Reagent Kit v2 Assay (Advanced Cell Diagnostics), following the manufacturer’s instructions with slight modifications. Briefly, hemocytes were pooled from five unfed ticks in microtubes containing 100 µL of L15C300. Then, cells were spun onto microscope slides using a Cytospin 2 and fixed with 4% paraformaldehyde (PFA, Millipore- Sigma) for 20 minutes. After three washing steps with 1X PBS, two drops of hydrogen peroxide were applied to the samples for 10 minutes and washed three times with 1X PBS. Subsequently, two drops of Protease III were added to the cells and the samples incubated in the HybEZ^TM^ Oven (Advanced Cell Diagnostics) for 30 minutes at 40°C. Samples were washed three times with 1X PBS before the addition of two drops of the following RNAScope probes (Advanced Cell Diagnostics): *hemocytin* RNA probe (regions 2778-3632 of XM_042287086.1) conjugated with C1, *astakine* probe (regions 47-800 of XM_040222953.1) conjugated with C2, *actin5c* probe (regions 2-1766 of XM_029977298.4) conjugated with C1 as a positive control and *gfp* probe (regions 12-686 of AF275953.1) conjugated with C1 as a negative control. Slides were incubated in a HybEZ^TM^ Oven for two hours at 40°C. Samples were then washed three times with 1X PBS and stored overnight in 5X SSC buffer (Millipore Sigma). Next, slides were washed using 1X PBS and incubated with the AMP reagents as instructed by the manufacturer. To develop the probe signals, samples were washed three times with 1X PBS and three drops of the corresponding RNAscope Multiplex FLv2 HRP was added for 15 minutes at 40°C. Then, slides were washed again with 1X PBS and incubated with ∼50 µL of 1:1500 dilution of Opal^TM^ 520 and 570 dyes for *hemocytin* and *astakine*, respectively (Akoya Biosciences) before the addition of two drops of the RNAscope Multiplex FLv2 HRP blocker for 15 minutes at 40°C. Slides were counterstained with two drops of DAPI for 30 seconds and mounted using two drops of ProLong™ Gold Antifade Mountant (Thermo Scientific). Slides were allowed to dry overnight before examined under a Nikon W-1 spinning disk confocal microscope. For imaging, the following laser channels were used: 488 nm (GFP, plasma membrane), and 405 nm (DAPI), 488 nm (GFP, C1) and 561 nm (RFP, C2).

### RNA interference

For *in vitro* experiments, small interfering RNAs (siRNAs) and scrambled controls (scRNAs) designed for *astikine* and *hemocytin* were synthesized by Millipore Sigma with UU overhangs. IDE12 cells were seeded at 3x10^5^ cells/well (24-well plate) for RNA extraction or the phagocytosis assay and 1x10^6^ cells/well (6-well plate) for protein extraction. siRNAs were transfected into IDE12 cells using Lipofectamine 3000 (Thermo Scientific) at 1 µg per mL. After 7 days, cells were either harvested or infected with *A. phagocytophilum* for 48 hours. For protein extraction, cells were washed with 1X PBS, resuspended in a solution of Radio- immunoprecipitation assay (RIPA) buffer (Merck Millipore) containing protease inhibitor cocktail (Roche) and stored at -80°C. For RNA extraction, cells were harvested in TRIzol and stored at - 80°C.

For *in vivo* experiments, siRNAs and scRNAs designed for *astakine* and *hemocytin* were synthesized using the Silencer™ siRNA Construction Kit (Thermo Scientific) with the primers listed in Supplementary Table 1, following the manufacturer’s instruction. Tick microinjections were performed using 40-60 ng of siRNA or scRNA per tick. Ticks were allowed to recover overnight before being placed on uninfected or *A. phagocytophilum-*infected mice. After placement, the percentage of ticks attached to mice was calculated. Fully fed ticks were collected 4 days post-placement, weighed, and either maintained in a humidified chamber for molting experiments or frozen at -80°C in 200 µL TRIzol for RNA extraction.

### Phagocytosis assay

IDE12 were transfected with siRNA or scRNA designed for *hemocytin,* as previously described (*7*). Briefly, 7 days post-transfection, cells were re-plated at a density of 4x10^5^ cells/well in a 24-well plate and left to adhere overnight. The following day, media was replaced with 500 μL of L15C300 containing a 1:10,000 dilution of FluoSpheres™ Carboxylate 1.0 µm beads (yellow-green, Invitrogen) for 24 hours. Cells were then washed five times with 1X PBS and fixed in 200 mL of 4% PFA for 20 minutes. Fluorescence and bright field microscopy images were acquired with an EVOS FL Digital Inverted Microscope (Advanced Microscopy Group) and merged using ImageJ software. Negative and positive cells for fluorescent microspheres and the average number of phagocytosed beads per cell were calculated from the number of green puncta detected in the total cells counted in each field (>100 cells). Two biological replicates were performed, with fields taken from three different wells per replicate.

### Recombinant *astakine*

*I. scapularis* recombinant Astakine (rAstk) was produced by GeneScript, based on the coding sequence of the XM_040222953.1 *astakine* transcript (LOC8032444). For *in vitro* cell proliferation assays, 1X10^5^ IDE12 cells/well were seeded in L15C300 complete medium (24-well plate). The following day, cells were treated with 0.05 µg/mL of rAstk for a total of 5 days. Cells were collected in a 1.5 mL microtube and 10 µL of the suspension was used for live cell counting using the trypan blue exclusion assay with a TC20™ automated cell counter (Bio-Rad). BSA (0.05 µg/mL) was used as negative control. For *in vivo* experiments, nymphs were microinjected with 50 nL of PBS containing 0.05 ng, 0.5 ng or 5 ng of rAstk (*27–29*). Afterwards, ticks were allowed to rest for 24 hours before hemolymph was collected, as described above. Microinjection of BSA (5 ng) was used as a negative control.

### Clodronate depletion of phagocytic hemocytes

To determine the effect of phagocytic hemocytes in *I. scapularis* nymphs, we used clodronate liposomes (CLD) to deplete phagocytic cells (*31, 36, 37*). Ticks were immobilized on a glass slide using double-sided tape and microinjected (Nanojet III microinjector; Drummond Scientific) through the anal pore with 69 nL of either CLD (Standard macrophage depletion kit, Encapsula NanoSciences LLC) or control liposomes (LP) at a 1:5 dilution in 1X PBS. Ticks were allowed to recover overnight before placement on uninfected mice. Following collection, fully fed ticks were weighed and frozen at -80°C in 200 µL of TRIzol for RNA extraction.

### RNA extraction and quantitative real-time PCR (qPCR)

Total RNA from cell lines or engorged ticks was isolated from samples preserved in TRIzol using the PureLink RNA Mini kit (Ambion). Total RNA from tick hemocytes was purified following the manufacturer’s instruction for TRIzol RNA extraction with the addition of 1 µg of glycogen (RNA grade, Thermo Scientific). For cDNA synthesis, 400-600 ng of RNA was used in the Verso cDNA Synthesis Kit (Thermo Scientific). To measure gene expression by qRT-PCR, 2X Universal SYBR Green Fast qPCR Mix (ABclonal) was used with the addition of 1 µL of cDNA per well and quantified using a CFX96 Touch Real-Time PCR Detection System (Bio-rad). All genes of interest were amplified at 54°C, except for amplification of *A. phagocytophilum* 16S, which was run at 46°C. No-template controls were included to verify the absence of primer-dimers formation and/or contamination. Reactions on each sample and controls were run in duplicate. Gene expression was determined by relative quantification normalized to tick *actin*, using the primers listed in Supplementary Table 1.

### Western blotting

To extract proteins, wells were washed with cold 1X PBS, and resuspended in 1X RIPA buffer (Merck Millipore) containing protease inhibitor cocktail (Roche). Protein extracts were stored at -80°C and their concentration was estimated using Bicinchoninic acid (BCA) assay (Thermo Scientific). For sodium dodecyl sulfate polyacrylamide gel electrophoresis (SDS-PAGE), equal amounts of protein were boiled in 6X Laemmli sample buffer (Alfa Aesar) containing 5% β-mercaptoethanol. The proteins were then transferred onto PVDF membranes (Biorad) and blocked for 1 hour using a solution of 5% skim milk prepared in 1X PBS and 0.1% Tween® 20 Detergent (PBS-T). Primary antibodies were incubated overnight at 4°C, followed by four washes in PBS-T and incubation with secondary antibodies for at least 1 hour at room temperature with gentle agitation. The blots were washed four times in PBST, incubated with enhanced chemiluminescence (ECL) substrate solution for 1 minute (Millipore), and then imaged. A complete list of antibodies used can be found in Supplementary Table 2.

### CRISPR activation (CRISPRa)

The SP-dCas9-VPR plasmid was a gift from George Church (Addgene plasmid, #63798).

For expression of sgRNAs, the sgRNA for the *hemocytin* promoter (or a scrambled version of its sequence) was cloned between the *BbsI* sites of a derivative of the pLib8 vector (pLibTB.1_ISCW_025) under the control of an *I. scapularis* U6 promoter (ISCW025025) (Supplementary Figure 16). ISE6 cells (7x10^7^ cells) were nucleofected with 15 µg of SP-dCas9- VPR plasmid using the buffer SF, pulse code EN150 (Lonza Biosciences) and subsequently incubated for 3 days. Following incubation, the cells were selected with neomycin (250 µg/mL) for 2 weeks. After antibiotic selection, the cells were nucleofected with 1 µg of the pLibTB.1_ISCW_025 vector containing the experimental or scrambled control sgRNA and were harvested 3 days later. Cells were collected in TRIzol for RNA extraction, as previously described.

### Statistics

Statistical significance for quantitative variables was tested with an unpaired t-test with Welch’s correction, Mann–Whitney U test or one-way analysis of variance (ANOVA) followed by Tukey’s multiple comparison test when appropriate. Gaussian distribution was assessed using the D’Agostino-Pearson omnibus K2 test, and homogeneity of variance was determined by F-test when comparing two conditions or the Brown-Forsythe test when comparing three conditions. Statistical significance for categorical variables was assessed using Fisher’s exact test. GraphPad PRISM® (version 9.1.2) was used for all statistical analyses. Outliers were detected by a Grubbs’ Test, using the GraphPad Quick Cals Outlier Calculator. P values of < 0.05 were considered statistically significant.

## Acknowledgments

The authors acknowledge members of the Pedra laboratory for providing insightful discussions and colleagues for manuscript feedback. We thank Jon Skare (Texas A&M University Health Science Center) for providing the *B. burgdorferi* B31 strain, clone MSK5; Ulrike G. Munderloh (University of Minnesota) for supplying ISE6 and IDE12 tick cells; Joseph Mauban (University of Maryland School of Medicine) for aiding in the microscopy analysis; Jonathan Oliver (University of Minnesota) for providing available *I. scapularis* ticks; Dana Shaw (Washington State University) and Adela Oliva Chavez (Texas A&M University) for information regarding cultures of *B. burgdorferi* and *A. phagocytophilum*; the University of Maryland, Baltimore Confocal Microscopy Core and the Maryland Genomics Core at the Institute for Genome Sciences for the services provided in microscopy and next generation sequencing, respectively. This work was supported by grants from the National Institutes of Health (NIH) to HJL-Y (T32AI162579), AJO (F31AI152215), LRB (F31AI167471), JHFP (R01AI134696, R01AI116523), JHFP and UP (P01AI138949), NP, SEM and JHFP (R21 AI168592). NP, SEM and JHF were also supported in-kind by the Fairbairn Family Lyme Research Initiative. NP is an investigator of the Howard Hughes Medical Institute. Some images were created with BioRender.com. The content is solely the responsibility of the authors and does not represent the official views of the NIH, the Department of Health and Human Services, NIST, the Department of Commerce or the United States government.

## Data and Code Availability

All sequencing reads for the bulk RNA sequences were deposited into the NCBI Sequence Read Archive under the BioProject accession PRJNA906572. All scRNA sequences were deposited into the NCBI Sequence Read Archive under the BioProject accession PRJNA905678. The codes for the scRNA sequence analysis are available through the GitHub websites: https://github.com/Haikelnb/scRNASeq_Analysis and https://github.com/HLaukaitisJ/PedraLab_hemocyte_scRNAseq. Reviewer tokens can be made available upon request.

## Resource Availability

Further information and request for resources and reagents should be directed to and will be honored by the corresponding author: Joao HF Pedra

## Author contributions

AR and JHFP designed the study. AR, HJL-Y, HNB, NS, SS, MB, CRF, AJO, MTM and LM performed the experiments. AR, HJL-Y and HNB performed computational analysis. MB, NS, EM and BX established the CRISPRa in the tick system. AR, HJL-Y, AJO and JHFP wrote the manuscript. LRB, LMV, VSR and FECP aided with experimentation. LRB created some schematics. All authors analyzed the data, provided intellectual input into the study, and contributed to editing of the manuscript. CS, SEM, UP, NP, and DS supervised experiments and provided instruments and reagents. JHFP supervised the study.

