## Supplementary Information for "Tick hemocytes have pleiotropic roles in microbial infection and arthropod fitness"

**Affiliations:** [1]Department of Microbiology and Immunology, University of Maryland School of Medicine, Baltimore, Maryland, USA.
[2]Institute for Genome Sciences, University of Maryland School of Medicine, Baltimore, Maryland, USA.
[3]Department of Genetics, Blavatnik Institute, Harvard Medical School, Boston, Massachusetts, USA.
[4]Howard Hughes Medical Institute, Chevy Chase, Maryland, USA.
[5]Department of Microbiology, National Emerging Infectious Diseases Laboratories, Boston University School of Medicine, Boston, Massachusetts, USA.
[6]Department of Veterinary Medicine, University of Maryland, College Park, Maryland, USA.
[†]Present address: Rancho BioSciences, San Diego, CA 92127
[#]Present address: Immunology Program, Memorial Sloan Kettering Cancer Center, New York, NY 10065

**This file includes:**
Supplementary Figures and Legends (S1 to S16)
Supplementary Tables (S1 to S2)
Supplementary Datasets (1 to 10)

### Supplementary Figure legends

**Supplementary Figure 1: Downregulated genes in hemocytes from engorged ticks compared to unfed.** The expression of (A) *osa*, (B) *runx1* and (C) *frizzled* was evaluated in hemocytes from unfed (ivory) or engorged (dark blue) ticks by RT-qPCR ($n$=6-11; samples represent 40-80 pooled ticks). Results represent mean ± SD. At least 6 biological replicates were performed. Statistical significance was evaluated by an unpaired t-test with Welch's correction. ***$p$<0.001; ****$p$<0.0001. *osa* = Brahma chromatin remodeling complex subunit osa; *runx1* = runt-related transcription factor 1; *frizzled* = frizzled-5.

**Supplementary Figure 2: Expression profile of immune genes identified through bulk RNA-seq.** Bar graph depicting fold change expression values of immune-related genes detected by bulk RNA-seq of hemocytes from engorged ticks compared to unfed ticks. Genes are color coded based on pathway or function.

**Supplementary Figure 3: Metrics of sequenced cells present in the hemolymph of *I. scapularis* nymphs.** Distribution and median of the **(A)** number (#) of unique molecular identifiers (UMIs), **(B)** genes and **(C)** percentage of mitochondrial transcripts (% mito) per cell in unfed (violet), uninfected engorged (teal), *A. phagocytophilum*-infected (pink) and *B. burgdorferi*-infected (green) *I. scapularis* hemocytes. The total number of cells sequenced per condition is depicted on the top of the graph. Low quality cells were filtered using the following thresholds: less than 600 unique reads, less than 150 expressed genes, and 30% or higher mitochondrial transcripts.

**Supplementary Figure 4: Principal component analysis (PCA) of sequenced cells present in the hemolymph of *I. scapularis* nymphs.** Variability between the unfed (red), uninfected engorged (green), *A. phagocytophilum*-infected (blue) and *B. burgdorferi*-infected (orange) hemocyte treatments is depicted.

**Supplementary Figure 5: t-Distributed Stochastic Neighbor Embedding (t-SNE) plot of cells collected from the hemolymph of unfed or engorged *I. scapularis* nymphs. (A)** Unfed (4,630 cells) and **(B)** engorged (15,802 cells) conditions. The engorged dataset contains samples from uninfected (6,000 cells), *A. phagocytophilum*-infected (6,287 cells) and *B. burgdorferi*-infected (3,515 cells) ticks.

**Supplementary Figure 6: Dot plots comparing the expression of marker genes from immune clusters between unfed and engorged ticks**. Marker genes of the **(A)** Immune 1 or **(B)** Immune 2 hemocyte cluster are represented in unfed (left) or engorged (right) ticks. The top 20 shared marker genes between conditions are shown. Average gene expression is demarked by intensity of color. Percentage of gene expression within individual clusters is represented by a dot diameter.

**Supplementary Figure 7: Dot plots comparing the expression of marker genes from proliferative clusters between unfed and engorged ticks.** Marker genes of the **(A)** Proliferative 1 or **(B)** Proliferative 2 hemocyte clusters are represented in unfed (left) or engorged (right) ticks. The top 20 shared marker genes between conditions are shown. Average gene expression is demarked by intensity of color. Percentage of gene expression within individual clusters is represented by a dot diameter.

**Supplementary Figure 8: Comparison of total hemocytes in uninfected and infected ticks.** Total hemocytes collected from *I. scapularis* nymphs fed on **(A)** *A. phagocytophilum*- (pink) or **(B)** *B. burgdorferi*- (green) infected mice compared to uninfected (blue; *n*=10-13). Results represent mean ± SD. At least two biological replicates were performed. Statistical significance was evaluated by an unpaired t-test with Welch's correction. (-) = Uninfected; Ap = *A. phagocytophilum*; Bb = *B. burgdorferi*. ns=not significant.

**Supplementary Figure 9: Attachment of RNAi-microinjected ticks. (A)** Attachment rate of *I. scapularis* nymphs microinjected with *hemocytin* siRNA (si-*hmc*) or scrambled RNA (sc-*hmc*) and placed on *A. phagocytophilum*-infected mice. **(B)** Attachment rate of *I. scapularis* nymphs microinjected with *astakine* siRNA (si-*astk*) or scrambled RNA (sc-*astk*) and placed on *A. phagocytophilum*-infected mice. Results are represented as percentage attached from total number of ticks placed. At least two biological replicates were performed. Statistical significance was evaluated by a Fisher exact test. ns = not significant.

**Supplementary Figure 10: *hemocytin* (*hmc*) expression in tick cells.** *hmc* expression in IDE12 relative to its abundance in ISE6 cells was evaluated by RT-qPCR (*n*=11-12). Results are represented as mean ± SD. Two biological replicates were performed. Statistical significance was evaluated by an unpaired t-test with Welch's correction. ****$p$<0.0001.

**Supplementary Figure 11: Hemocytes in *hemocytin* (*hmc*)-silenced ticks. (A)** Total number of hemocytes and **(B)** percentage of morphotypes in ticks microinjected with *hmc* siRNA (si-*hmc*; blue) or scrambled RNA (sc-*hmc*; grey) fed on uninfected mice (*n*=9-11). Results are represented as mean ± SD. At least two biological replicates were performed. Statistical significance was evaluated by an unpaired t-test with Welch's correction. ns=not significant.

**Supplementary Figure 12: Phagocytosis in *hemocytin* (*hmc*)-silenced tick cells.** IDE12 cells were transfected with *hmc* siRNA (si-*hmc*) or scrambled RNA (sc-*hmc*) for seven days prior to adding fluorescent microspheres for 24 hours. Data were quantified as **(A)** percentage of cells positive for fluorescent beads or **(B)** the average number of beads per cell in each field. 109-115 cells were analyzed per replicate (*n*=3). Data represent two combined independent experiments. Bead+ = cells positive for beads; Bead- = cells negative for beads. Statistical significance was evaluated by an unpaired t-test. ns= not significant.

**Supplementary Figure 13: Proliferation in IDE12 cells treated with recombinant Astakine.** 0.05 µg/mL of rAstk or BSA (control) were added to $1 \times 10^5$ IDE12 cells for 5 days before total number of live cells were counted (*n*=17). Results are represented as mean ± SD. Three biological replicates were performed. Statistical significance was evaluated by an unpaired t-test with Welch's correction. **$p<0.01$. rAstk = recombinant Astakine; BSA = Bovine Serum Albumin.

**Supplementary Figure 14: Attachment of microinjected ticks to mice.** Attachment rate of *I. scapularis* nymphs microinjected with **(A)** clodronate or empty liposomes (control), **(B)** *hemocytin* siRNA (si-*hmc*) or scramble RNA (sc-*hmc*) or **(C)** *astakine* siRNA (si-*astk*) or scramble RNA (sc-*astk*) and placed on uninfected mice. Results present percentage of ticks attached out of total number placed. At least two biological replicates were performed. Statistical significance was evaluated by a Fisher exact test. ns = not significant.

**Supplementary Figure 15: Weight of microinjected ticks after a blood meal. (A)** Weight of engorged nymphs microinjected with *hemocytin* siRNA (si-*hmc*; blue) or scramble RNA (sc-

*hmc*; grey) fed on *A. phagocytophilum*-infected mice ($n$=11-14). **(B)** Weight of engorged nymphs microinjected with *astakine* siRNA (si-*astk*; blue) or scrambled RNA (sc-*astk*; grey) fed on *A. phagocytophilum*-infected mice ($n$=15-18). Results are represented as mean ± SD. Two biological replicates were performed. Statistical significance was evaluated by an unpaired *t*-test with Welch's correction. $**p<0.01$.

**Supplementary Figure 16: Engineering of the library vector (pLibTB.1_ISCW_025) for single guide (sg) RNA expression.** (Left) Cartoon representing the pLibTB.1_ISCW_025 vector used for sgRNA cloning. The 20 bp sgRNA targeting the first exon of the *hemocytin* (*hmc*) gene (ISCP_023268) or scrambled control was cloned between *BbsI* sites. The plasmid was constructed using the *I. scapularis* U6 promoter sequence for optimal expression of the sgRNA. (Right) Sequence of the *I. scapularis* U6 promoter and the sgRNA targeting *hemocytin*: *I. scapularis* U6 promoter (ISCW025025; blue); sgRNA spacer for *hmc* or scramble (purple); gRNA scaffold (green); 8-T terminator (black) and U6 snRNA downstream region (3'-ISCW025025; orange). CAG promoter includes: Cytomegalovirus (CMV) early enhancer element; promoter, first exon and first intron of the chicken *β-actin* gene and splice acceptor of the rabbit *β-globin* gene. EBFP2: constitutively fluorescent enhanced blue fluorescent protein; T2A: *Thosea asigna* virus ribosome skipping sequence; PuroR: puromycin resistance (puromycin N-acetyl transferase); SV40: Simian virus 40 termination and poly-adenylation sequence; AmpR: ampicillin resistance (TEM-1 β-lactamase); Ori: origin of replication; attB: phi-C31 attB sites for recombinase-mediated cassette exchange.

**Supplementary Table 1:** Primers and siRNA sequences

| Target | Type | Start Position Within mRNA | siRNA/ Primer Name | Strand | Primer Sequence | Accession number |
|---|---|---|---|---|---|---|
| ***I. scapularis hemocytin*** | siRNA | 6891 | siHmc_F | forward | AAGCCAATAAGTCAGATGGT TCCTGTCTC | XM_042287086.1 |
| | | | siHmc_R | reverse | AAAACCATCTGACTTATTGG CCCTGTCTC | |
| | scrambled | | scHmc_F | forward | AAGTCTGAGCACGTTAATAG ACCTGTCTC | |
| | | | scHmc_R | reverse | AAGTCTGAGCACGTTAATAG ACCTGTCTC | |
| | qRT-PCR | 4690 | Hmc_F | forward | AATGGCACCATCACAGAGGA | |
| | | 4756 | Hmc_R | reverse | GTGAAGTTCCTGACGCACTC | |
| ***I. scapularis astakine*** | siRNA | 623 | siAstk_F | forward | AACCAAGGACGATTTCGGCA ACCTGTCTC | XM_040222953.1 |
| | | | siAstk_R | reverse | AATTGCCGAAATCGTCCTTG GCCTGTCTC | |
| | scrambled | | scAstk_F | forward | AAGGAACGTATCGGCATCAC ACCTGTCTC | |
| | | | scAstk_R | reverse | AATGTGATGCCGATACGTTC CCCTGTCTC | |
| | qRT-PCR | 92 | Astk_F | forward | CAGCGTAGCGTTAGTAGCAC | |
| | | 233 | Astk_R | reverse | GTTCGATGGGCAGAGAATG G | |
| ***I. scapularis jun*** | qRT-PCR | 332 | Jun_F$ | forward | CGCAGTACCTGTTCACGAAG | |
| | | 497 | Jun_R$ | reverse | CGACGACGAGGGTAAGATG A | |
| ***I. scapularis frizzled-5*** | qRT-PCR | 4477 | Frizz_F | forward | CGTGATCTGTGGTCCGTCTA | XM_029968494.4 |
| | | 4595 | Frizz_R | reverse | ATCCGGTGAAACCTGGGAAT | |

| | | | | | | |
|---|---|---|---|---|---|---|
| ***I. scapularis runt-related transcription factor 1*** | qRT-PCR | 2063 | Runx1_F | forward | TGTTGGATGACTCCGGTTGA | XM_029992700.4 |
| | | 2176 | Runx1_R | reverse | AGCTCCTTCGGTCTTTCCAA | |
| ***I. scapularis trithorax group protein osa*** | qRT-PCR | 6677 | Osa_F | forward | AAACCGATCCCTCGGAGTTG | XM_040206018.3 |
| | | 6772 | Osa_R | reverse | ACTGCTCAAATTGCCAAGCA | |
| ***I. scapularis jnk*** | qRT-PCR | 254 | Jnk_F | forward | GGATCACGTCAGCAGGTCTA | XM_002408785.5 |
| | | 417 | JNK_R | reverse | ATTGTGACCAGTCACGGAGT | |
| ***I. scapularis glucose-6-phosphatase 2*** | qRT-PCR | 1486 | G6Pase2_F | forward | ACCAACGTGCCTGTCTTCTA | XM_029991048.3 |
| | | 1605 | G6Pase2_R | reverse | TGGAACCGGCTTAGTTGGAT | |
| ***I. scapularis inosine-5'-monophosphate dehydrogenase 1*** | qRT-PCR | 817 | IMPDH1_F | forward | GTACAAGCACGGGTTCATCC | XM_029980674.4 |
| | | 960 | IMPDH1_R | reverse | GATGTCCCTTGAGGTCACCA | |
| ***I. scapularis 4-coumarate-CoA ligase 1*** | qRT-PCR | 489 | 4CL1_F | forward | TCCGAGCGTAATGAAAGGGT | XM_029976738.4 |
| | | 619 | 4CL1_R | reverse | CAGCTGATCACTTCGTGCAA | |
| ***I. scapularis adenosylhomocysteinase B*** | qRT-PCR | 618 | AHCY_F | forward | GTCATGATTGCTGGCAAGGT | XM_029975123.4 |
| | | 705 | AHCY_R | reverse | ATCTCCGTAACAAGCACCCT | |
| ***I. scapularis croquemort*** | qRT-PCR | 578 | Crq_F$ | forward | CCCAAGTTGCAGGAGGTGG | XM_029984267.4$ |
| | | 663 | Crq_R$ | reverse | CGCACCTCCCGGTAGG | |
| ***A. phagocytophilum16S*** | qRT-PCR | 116 | Ap 16S_F† | forward | GGTGAGTAATGCATAGGAATC | APH_RS03965# |
| | | 223 | Ap 16S_R† | reverse | GCTCATCTAATAGCGATAAATC | |
| ***B. burgdorferi recA*** | qRT-PCR | 194 | RecA_F# | forward | GTGGATCTATTGTATTAGATGAGGCT | U23457.1 |

| | | | | | | |
|---|---|---|---|---|---|---|
| | | 390 | RecA_R# | reverse | GCCAAAGTTCTGCAACATTAACACCT | |
| ***I. scapularis actin*** | qRT-PCR | 896 | I.s. actin_F† | forward | GGTCATCACAATCGGCAAC | XM_029977298 |
| | | 1003 | I.s. actin_R† | reverse | ATGGAGTTGTACGTGGTCTC | |
| ***hmc* sgRNA** | sgRNA | +20 bp from ATG | | | CTTGTCCCGGTTTCTGCAGCCGG | ISCP_023268 |
| **sc sgRNA** | sgRNA | scramble control | | | GGAGCAATAGGACCGTGCTGAGG | N/A |

#Shaw *et al.*, 2017; DOI:10.1038/ncomms14401
†Oliva Chavez *et al.*, 2021; DOI:10.1038/s41467-021-23900-8
$O'Neal *et al.*, 2023; DOI:10.1073/pnas.2208673120

**Supplementary Table 2**: Resources and reagents available

| **Antibody** | **Source** | **Identifier** | **Dilution/ Concentration** |
|---|---|---|---|
| Goat Anti-mouse IgG H+L(HRP) | Elabscience | E-AB-1001 | 1:3,000 |
| Goat Anti-Rabbit IgG H&L (HRP) | Abcam | ab97051 | 1:4,000-10,000 |
| Rabbit Anti-Mouse Actin | Millipore Sigma | A2103 | 1:4,000 |
| Rabbit Phospho-SAPK/JNK (Thr183/Tyr185) Polyclonal Ab | Cell Signaling | 4668 | 1:1,000 |
| Rabbit JNK Polyclonal Ab | Proteintech | 10023-1-AP | 1:1,000 |
| Mouse Anti-*I. scapularis* Relish monoclonal Ab | GenScript | custom | 1:500 |
| **Cell Dye** | | | |
| Opal™ 520 dye | Akoya Biosciences | FP1487001KT | 1:1,500 |
| Opal™ 570 dye | Akoya Biosciences | FP1488001KT | 1:1,500 |
| ProLong™ Gold Antifade Mountant | Thermo Scientific | P10144 | N/A |
| **Cell Media** | | | |
| Leibovitz's L-15 Medium, powder | Gibco | 41300039 | N/A |
| L-aspartic acid | Millipore-Sigma | 11189 | 0.449 g/L |
| L-glutamine | Millipore-Sigma | G8540 | 0.500 g/L |
| L-proline | Millipore-Sigma | 81709 | 0.450 g/L |
| L-glutamic acid | Millipore-Sigma | 49449 | 0.250 g/L |
| alpha-ketoglutaric acid | Millipore-Sigma | K1128 | 0.449 g/L |
| Sodium hydroxide | Millipore-Sigma | S8045 | 10 N |
| D-glucose | Millipore-Sigma | G7021 | 18.018 g/L |
| FBS (USDA approved; for tick media) | Millipore-Sigma | F0926-500ML | 0.1 |
| Bacto™ Tryptose Phophate Broth | BD | 260300 | 0.1 |
| Lipoprotein Concentrate | MP Biomedicals | 191476 | 0.001 |

| | | | |
|---|---|---|---|
| Normal rabbit serum | Pel-Freez | 31126-5 | 0.06 |
| Sodium bicarbonate | Millipore-Sigma | S6014 | 0.0025 |
| HEPES | Millipore-Sigma | H4034 | 25 mM |
| CMRL1066 w/L-Glutamine (Powder) | US Biological | C5900 | N/A |
| Sodium citrate tribasic dihydrate | Millipore-Sigma | S4641 | 0.7 g/L |
| Yeastolate | BD | 255772 | 2 g/L |
| Neopeptone | BD | 211681 | 5 g/L |
| RPMI-1640 Medium With L-Glutamine | Quality Biological | 112-025-101 | N/A |
| Fetal Bovine Serum (for HL-60s media) | Gemini Bio-Products | 100-106 | 0.1 |
| GlutaMax | Gibco | 35050-061 | 0.01 |
| Rifampicin | Millipore-Sigma | 557303 | 50 mg/ml |
| Phosphomycin | Millipore-Sigma | P5396 | 100 mg/ml |
| Amphotericin B | Gibco | 15290-026 | 1:100 |
| Sodium pyruvate | Millipore-Sigma | P5280 | 0.8 g/L |
| Distilled water | Gibco | 15-230-147 | N/A |
| N-Acetyl-α-D-glucosamine | Millipore-Sigma | 1079-25GM | N/A |
| Albumin, Bovine Fraction V | MP Biomedicals | 160069 | N/A |
| Ampicillin | Millipore-Sigma | A0166 | 100 mg/ml |
| **Materials** | | | |
| Cellstar® cell culture flasks, 25 $cm^2$ | Greiner bio-one | 690-160 | N/A |
| T-25 Vented Flasks | CytoOne | CC7682-4825 | N/A |
| Cell culture plate with lid (6 well, flat bottom) | Millipore-Sigma | SIAL0516 | N/A |
| Costar® cell culture plate with lid (24 well, flat bottom) | Corning | CLS3526-1EA | N/A |
| NuPAGE™ 4-12% Bis-Tris Protein Gels, 1.5 mm, 10-well | Thermo Scientific | NP0335BOX | N/A |

| | | | |
|---|---|---|---|
| Mini-Protean® TGX™ gels | Biorad | 456-9034 | N/A |
| Trans-blot® Turbo™ Tranfer pack, 0.2 µm PVDF | Biorad | 1704156 | N/A |
| 5% Mini-PROTEAN® TBE Gel | Biorad | 4565014 | N/A |
| Biodyne B Precut Nylon Membranes | Thermo Scientific | 77016 | N/A |
| FALCON® 14 ml Polypropylene round-bottom tube | Corning | 352059 | N/A |
| 250 mm glass desiccator | Thermo Scientific | 08-615B | N/A |
| 1.5 ml microcentrifuge tubes | Thermo Scientific | 1148T71 | N/A |
| 15 ml conical screw cap tubes | USA Scientific | 5618-8261 | N/A |
| 50 ml conical screw cap tubes | USA Scientific | 5622-7270 | N/A |
| 500 ml vacuum filter/storage bottle system, 0.2 µm | Corning | 430773 | N/A |
| Rnase-free disposable pellet pestles | Thermo Scientific | 12-141-368 | N/A |
| 27 gauge, 1/2" needle | BD | 305109 | N/A |
| Siliconized tips | VWR | 53503-800 | N/A |
| Microcentrifuge Tubes with Socket Screw-Cap | VWR | 89004-304 | N/A |
| Non-stick RNAase-free 1.5 mL microtubes | Ambion | AM12450 | N/A |
| Fisherbrand™ Superfrost™ Plus Microscope Slides | Thermo Scientific | 12-550-15 | N/A |
| Double Sided Tape Mounting Tape Heavy Duty, 164 FT Length, 0.31 Inch/ 8mm Width | 3M | B093K3S2QL | N/A |
| White Filter Paper for CytoSep™ Single Funnel | Simport Scientific | M965FW | N/A |
| Flowmi® 40 uM Cell Strainers | Millipore-Sigma | BAH136800040 | N/A |
| **Reagents** | | | |
| Halt™ phosphatase inhibitor cocktail (100x) | Thermo Scientific | 78426 | 0.111111111 |
| Halt™ protease inhibitor cocktail (100x) | Thermo Scientific | 87786 | 0.111111111 |
| 10X RIPA | Millipore-Sigma | 20-188 | 1X |

| | | | |
|---|---|---|---|
| Chloroform | Millipore-Sigma | 288306 | 1 |
| Ethyl alcohol, Pure; 200 proof for molecular biology | Millipore-Sigma | E7023-1L | 70-100% |
| Lipofectamine 3000 Reagent | Thermo Scientific | L3000008 | 7.5 ml/1 ml |
| TRIzol reagent | Ambion | 15596018 | N/A |
| 2X Universal SYBR Green Fast qPCR Mix | Abclonal | RK21203 | 1X |
| Isoflurane, USP | MWI Veterinary Supply Co | 13985-528-60 | 3% |
| JumpStart™ REDTaq® ReadyMix™ Reaction Mix | Millipore Sigma | P1107 | 1X |
| 1X PBS | Quality Biological | 114-058-101 | N/A |
| OptiPrep™ Density Gradient Medium | Millipore-Sigma | D1556-250ML | 1.09 g/mL in PBS |
| RNasin® Plus Ribonuclease Inhibitor | Promega | N2611 | 0.2 U |
| HyClone™ Water, Molecular Biology Grade | Cytiva | SH3053801 | N/A |
| Sodium Chloride | Millipore-Sigma | S7653 | 300 mM |
| 2-βmercaptoethanol | Gibco | 21985-023 | 0.05 |
| 6X Laemmli buffer | Alfa Aesar | J60660 | 1X |
| Blocking grade blocker, non fat skim milk | Biorad | 1706404 | 0.05 |
| Bovine Serum Albumin | Millipore-Sigma | A2058 | 0.03 |
| Sodium dodecyl sulfate (SDS) | Millipore-Sigma | L6026 | 0.001 |
| Paraformaldehyde | Millipore-Sigma | P6148 | 0.04 |
| Immobilon Forte Western HRP substrate | Millipore-Sigma | WBLUF0100 | N/A |
| SSC Buffer 20× Concentrate | Millipore-Sigma | S6639 | 5X |
| SPRIselect for Size Selection | Beckman Coulter Genomics | B23319 | |
| FluoSpheresCarboxylate-Modified Microspheres | Thermo Scientific | F8823 | 1:10,000 |
| RNAscope™ Probe- Is-LOC8025528-C1 | Advanced Cell Diagnostics | 1215511-C1 | 2 drops / slide |
| RNAscope™ Probe- Is-LOC8032444-C2 | Advanced Cell Diagnostics | 1215521-C2 | 2 drops / slide |
| RNAscope™ Probe- Is-LOC8027393-C1 | Advanced Cell Diagnostics | 1164711-C1 | 2 drops / slide |
| RNAscope™ Probe- GFP | Advanced Cell Diagnostics | 409011 | 2 drops / slide |

| | | | |
|---|---|---|---|
| Glycogen, RNA grade | Thermo Scientific | R0551 | 1 mg |
| BbsI-HF | New England Biolabs | R3539S | 10 u/mg of DNA |
| Gum rosin | Millipore-Sigma | 60895 | 75% (3/4 parts) |
| Beeswax | Thermo Scientific | S25192A | 25% (1/4 parts) |
| **Commercial Assays** | | | |
| Richard-Allan Scientific™ Three-Step Stain Set | Thermo Scientific | 3300 | N/A |
| Pure Link RNA mini kit | Ambion | 12183025 | N/A |
| Silencer™ siRNA Construction Kit | Thermo Scientific | AM1620 | N/A |
| Verso cDNA Synthesis Kit | Thermo Scientific | AB-1453B | N/A |
| RNAscope® Multiplex Fluorescent Reagent Kit V2 | Advanced Cell Diagnostics | 323100 | N/A |
| Pierce BCA Protein Assay Kit | Thermo Scientific | 23227 | N/A |
| Pierce ECL Western Blotting Substrate | Thermo Scientific | 32106 | N/A |
| Standard Macrophage Depletion Kit (Clodrosome® + Encapsome®) | Encapsula NanoSciences LLC | CLD-8901 | 1:5 in PBS |
| Endofree Plasmid Maxi Kit | Qiagen | QGN-12362 | N/A |
| QIAamp® DNA Mini Kit | Qiagen | 51304 | N/A |
| QIAprep® Spin Miniprep Kit | Qiagen | 27106 | N/A |
| *Mycoplasma* testing kit | Southern Biotech | 13100-01 | N/A |
| SF Cell Line 4D-NucleofectorTM X Kit L | Lonza Bioscience | V4XC-2012 | N/A |
| NEBNext® Ultra™ II Directional RNA Library Prep Kit for Illumina® | New England Biolabs | E7760S | N/A |
| **Equipment** | | | |
| 4D-Nucleofector™ System | Lonza Bioscience | AAF-1002 | N/A |
| Nanoject III | Drummond Scientific Company | 3-000-207 | N/A |
| Epredia™ Cytospin™ 4 Cytocentrifuge | Thermo Scientific | A78300003 | N/A |

| | | | |
|---|---|---|---|
| CFX96 Touch Real-Time PCR Detection System | Biorad | Discontinued | N/A |
| C1000 Touch Thermocycler | Biorad | 1851148 | N/A |
| Percival I30BLL incubator | Percival | I30BLL | N/A |
| Vannas Spring Scissors - 4mm Cutting Edge | Fine Science Tools | 15018-10 | N/A |
| HybEZ™ II Oven | Advanced Cell Diagnostics | 321719 | N/A |
| Carl Zeiss™ Primo Star™ HAL/LED Microscope | Carl Zeiss | 415500-0057-000 | N/A |
| Axiocam 305 color camera | Carl Zeiss | 426560-9030-000 | |
| **Plasmids** | | | |
| SP-dCas9-VPR | Addgene | 63798 | N/A |
| pLibTB.1_ISCW_025 | Perrimon laboratory | N/A | N/A |
| **Cell Lines** | | | |
| *I. scapularis* ISE6 cells | Ulrike Munderloh, University of Minnesota | ISE6 | N/A |
| *I. scapularis* IDE12 cells | Ulrike Munderloh, University of Minnesota | IDE12 | N/A |
| HL-60 | ATCC | CCL-240 | N/A |
| **Organisms** | | | |
| *I. scapularis* nymph ticks | Tick Lab, Oklahoma State University | N/A | N/A |
| *I. scapularis* nymph ticks | Ulrike Munderloh, University of Minnesota | N/A | N/A |
| C57BL6J mice | University of Maryland, Baltimore | N/A | N/A |
| C57BL6J mice | Jackson Laboratories | #000664 | N/A |
| C3H/HeJ mice | Jackson Laboratories | #000659 | N/A |
| *A. phagocytophilum* HZ | Ulrike Munderloh, University of Minnesota | N/A | N/A |

| | | | |
|---|---|---|---|
| *B. burgdoferi* B31 clone MSK5 | Jon Skare, Texas A&M University Health Science Center | N/A | N/A |
| One Shot Top10 Chemically Competent E. Coli | Thermo Scientific | C404003 | N/A |
| **Software** | | | |
| VectorBase v63 | https://vectorbase.org/vectorbase/app | N/A | N/A |
| FlyBase vFB2023_02 | https://flybase.org/ | N/A | N/A |
| InterPro | https://www.ebi.ac.uk/interpro/ | N/A | N/A |
| UniProt v2023_02 | https://www.uniprot.org/ | N/A | N/A |
| Single-Cell RNA-seq data portal of DRSC/Perrimon lab | https://www.flyrnai.org/scRNA/blood/ | N/A | N/A |
| ImageJ v1.53t | https://imagej.nih.gov/ij/ | N/A | N/A |
| GraphPad Prism v9.1.2 | https://www.graphpad.com/ | N/A | N/A |
| Graphpad Quick Cals Outlier Calculator | https://www.graphpad.com/quickcalcs/Grubbs1.cfm | N/A | N/A |
| ZEN v3.6.095.08000 | https://www.micro-shop.zeiss.com/en/us/softwarefinder/software-categories/zen-blue/zen-blue-pro/ | N/A | N/A |

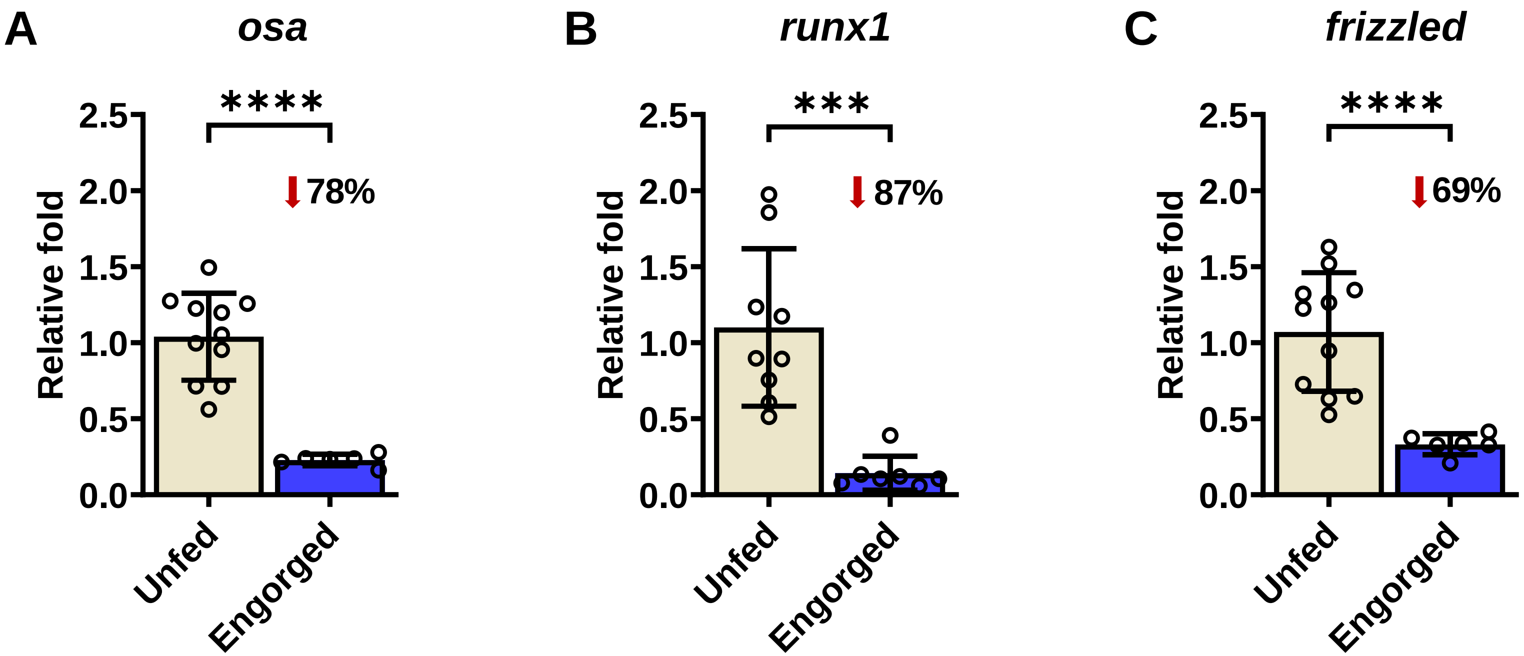

**Rolandelli *et al.* - Suppl. Fig. 1**

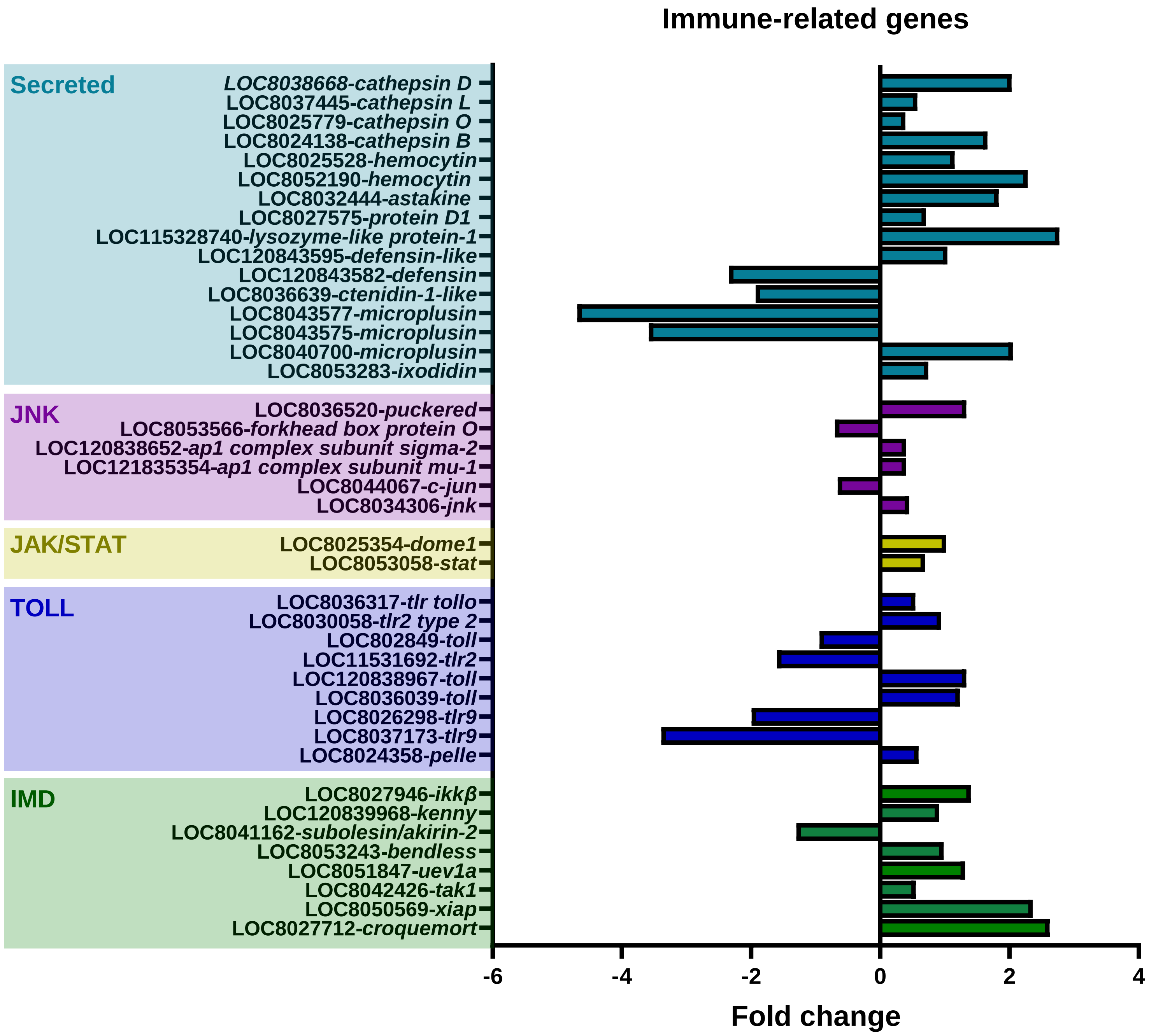

Rolandelli *et al.* - Suppl. Fig. 2

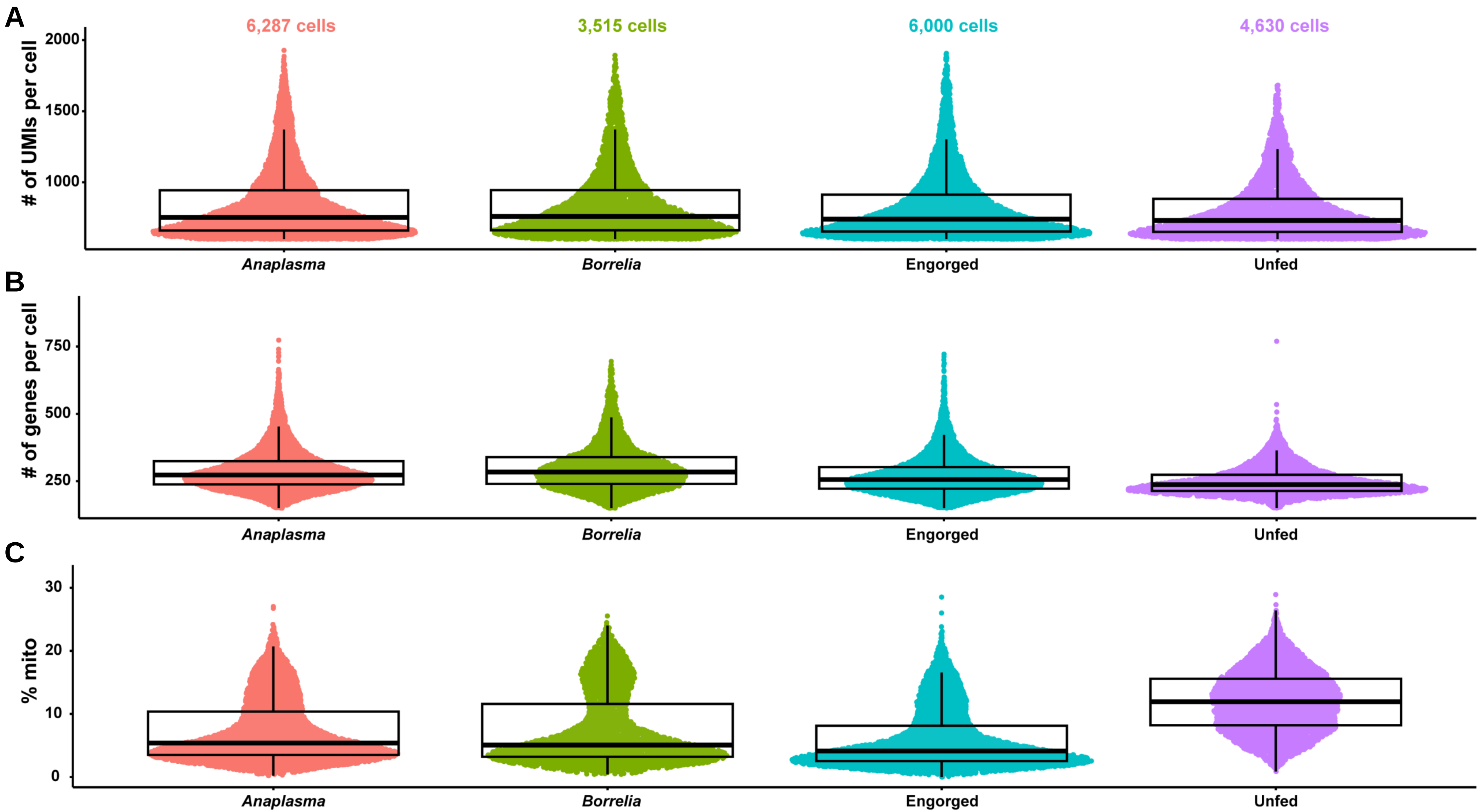

Rolandelli *et al.* - Suppl. Fig. 3

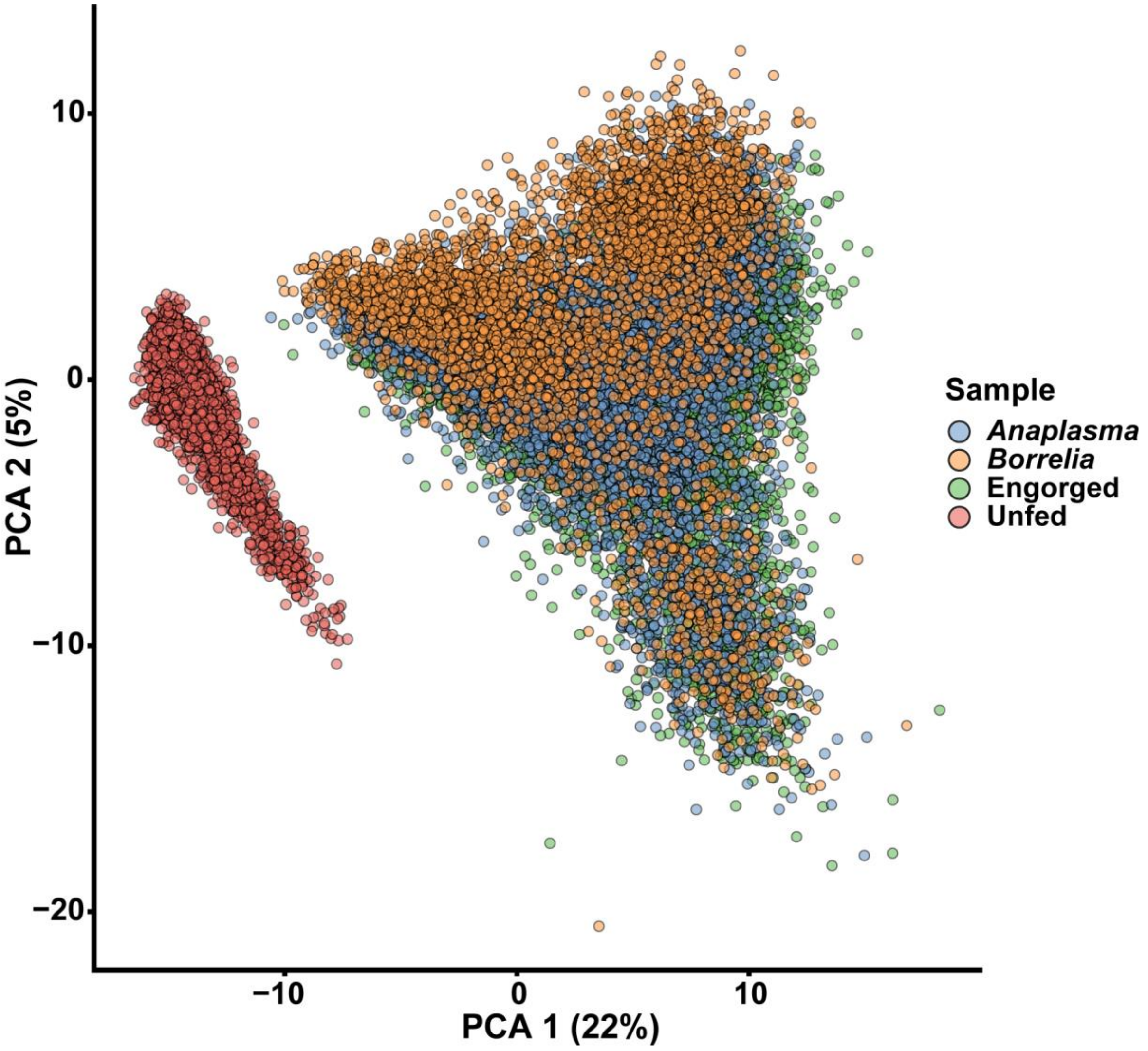

**Rolandelli *et al.* - Suppl. Fig. 4**

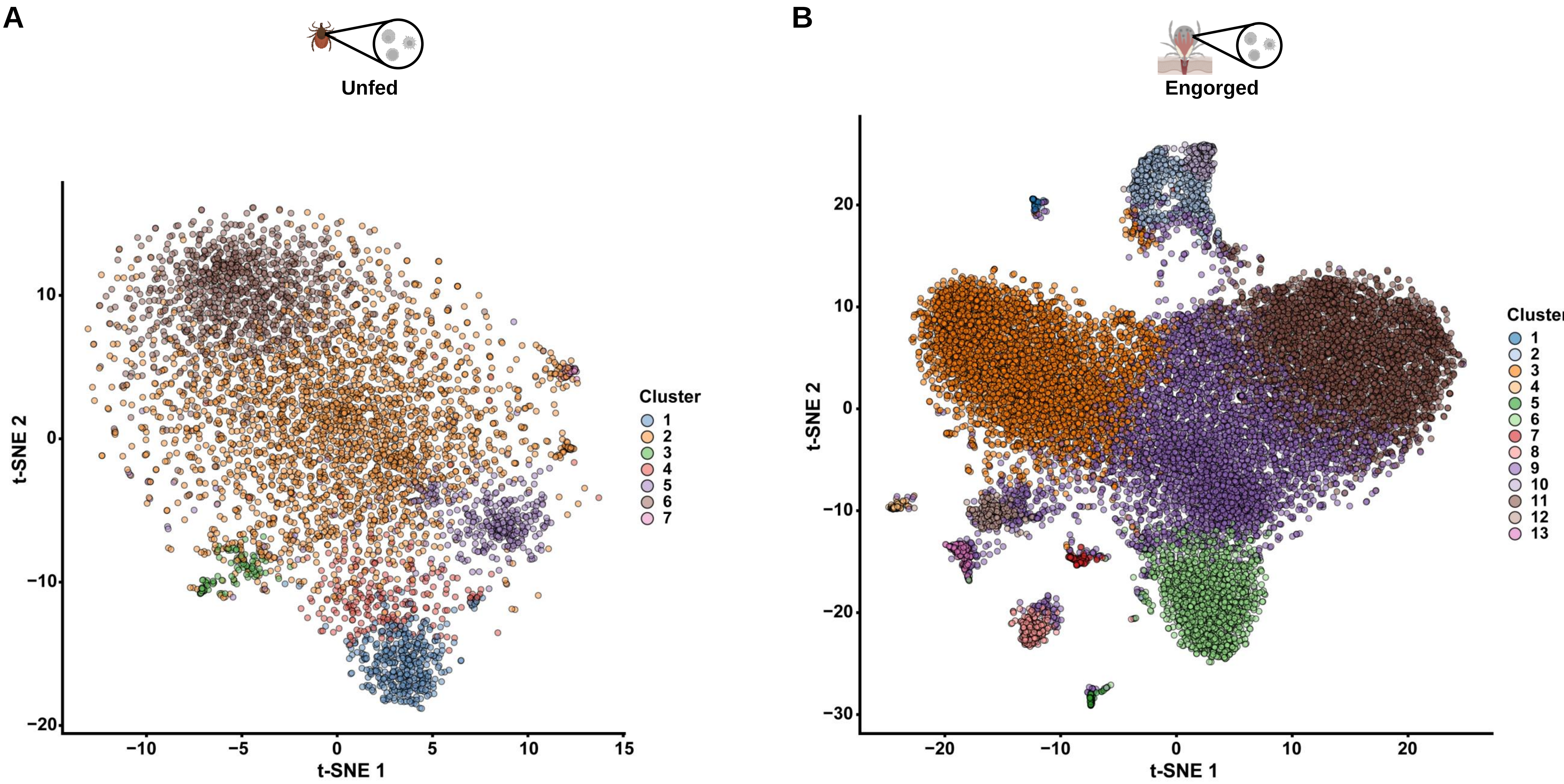

Rolandelli *et al.* - Suppl. Fig. 5

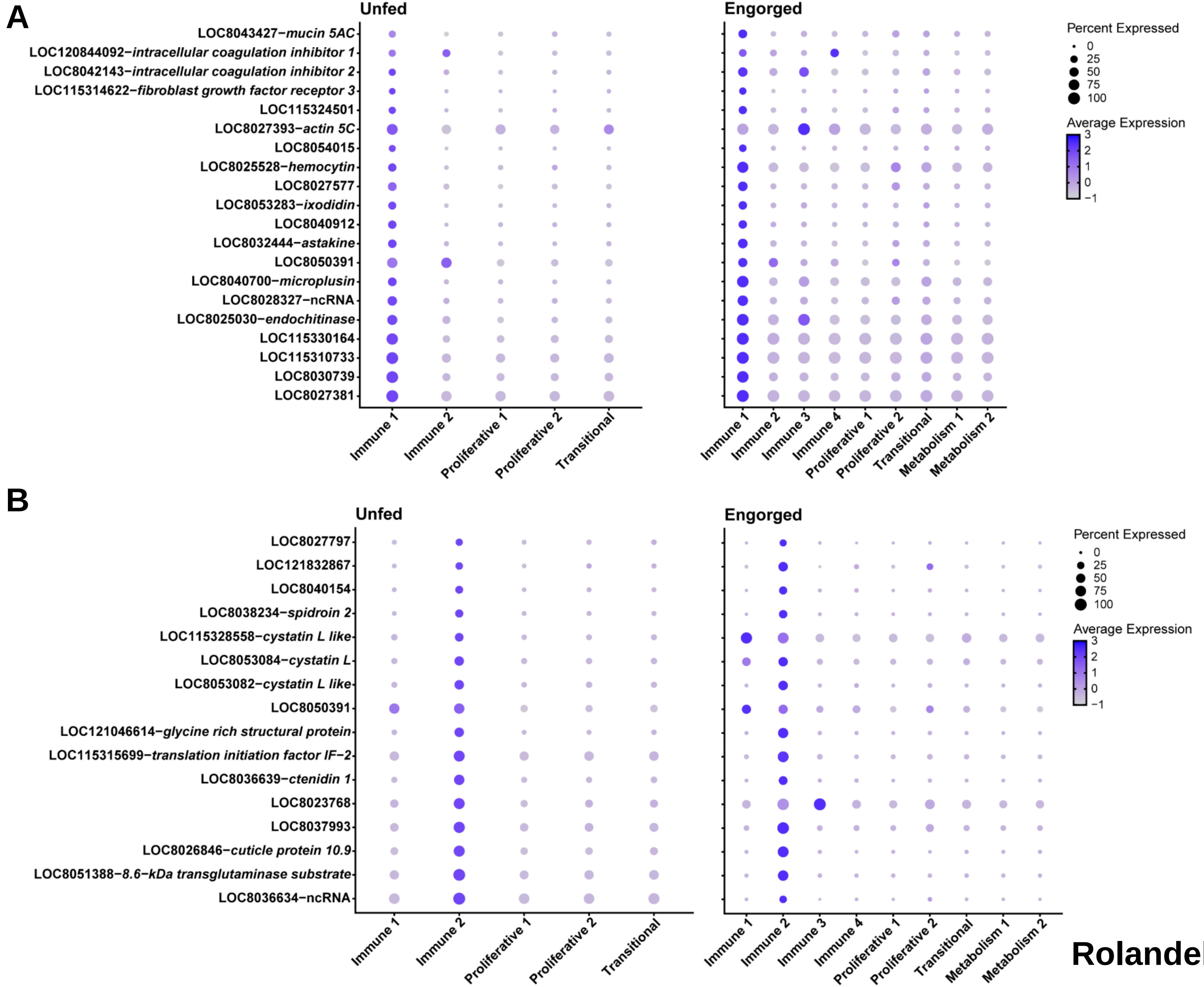

Rolandelli *et al.* - Suppl. Fig. 6

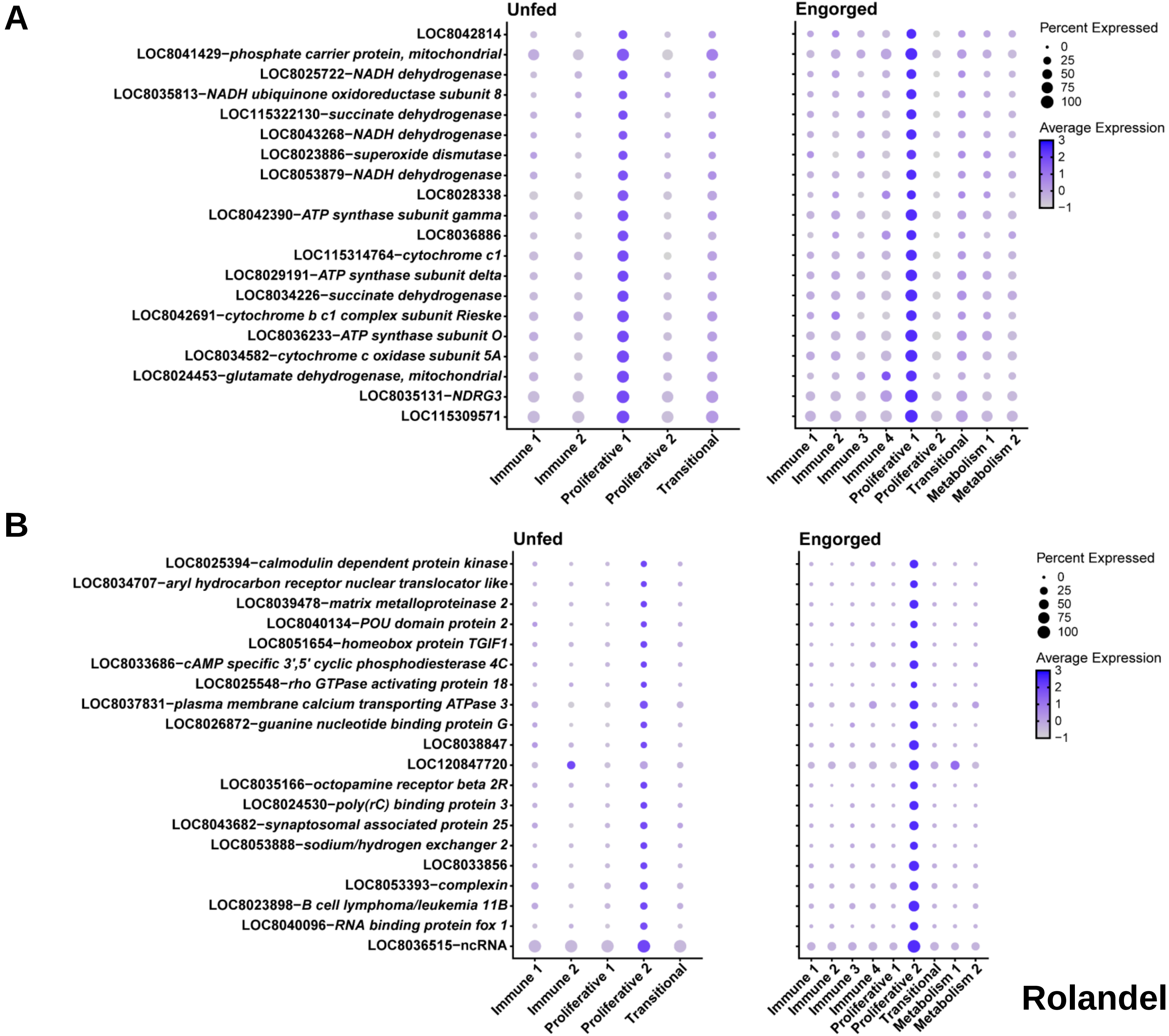

**Rolandelli *et al.* - Suppl. Fig. 7**

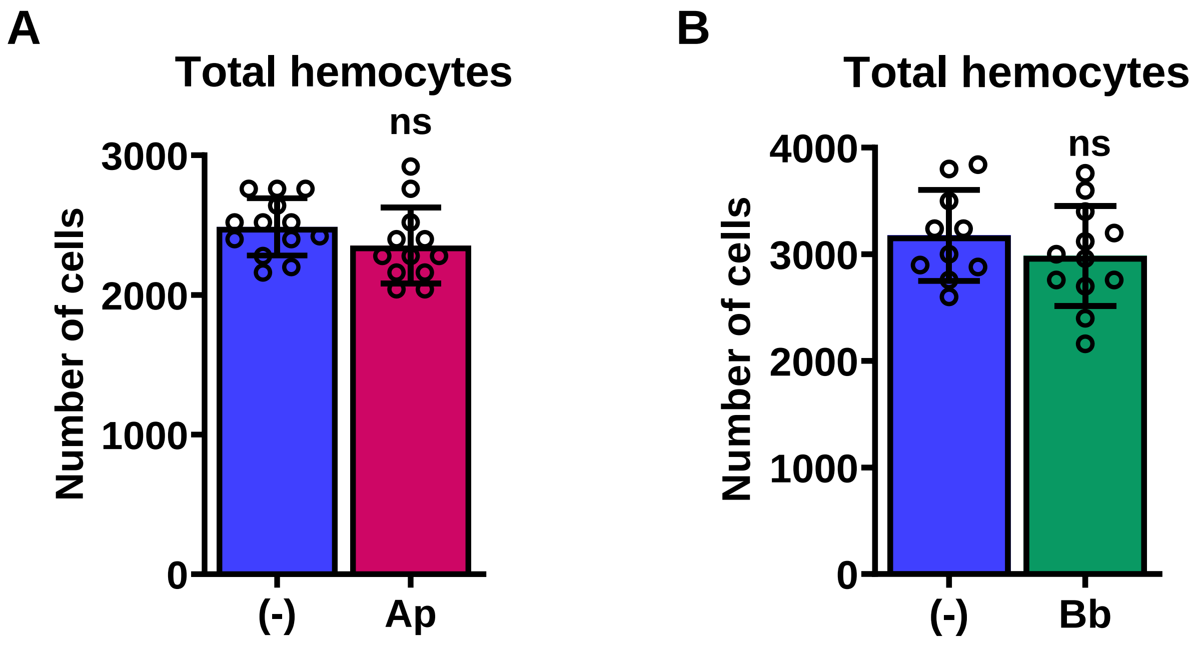

**Rolandelli *et al.* - Suppl. Fig. 8**

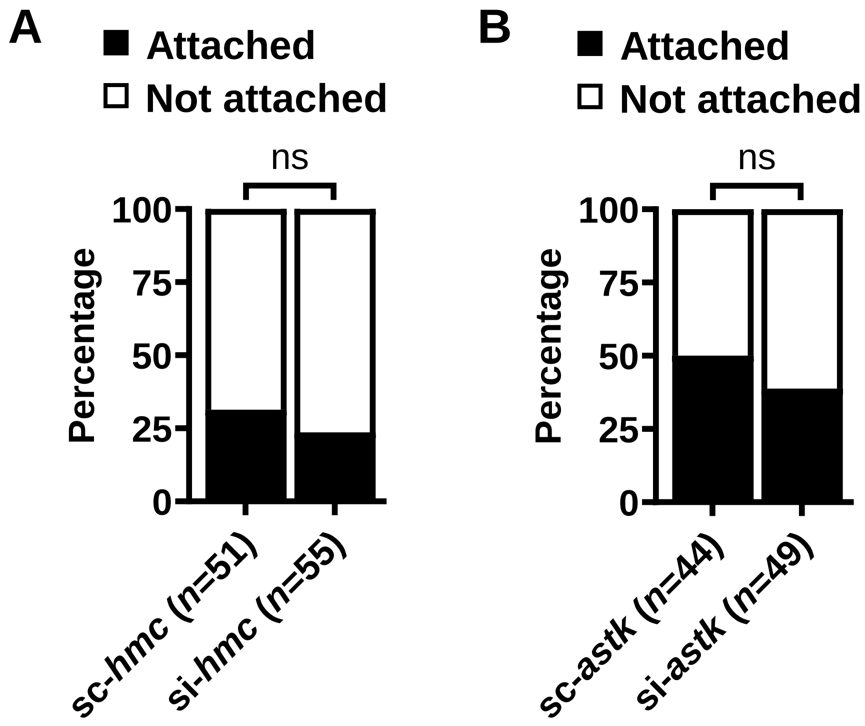

Rolandelli *et al.* - Suppl. Fig. 9

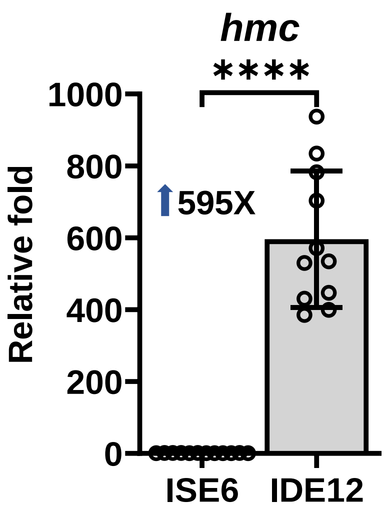

Rolandelli *et al.* - Suppl. Fig. 10

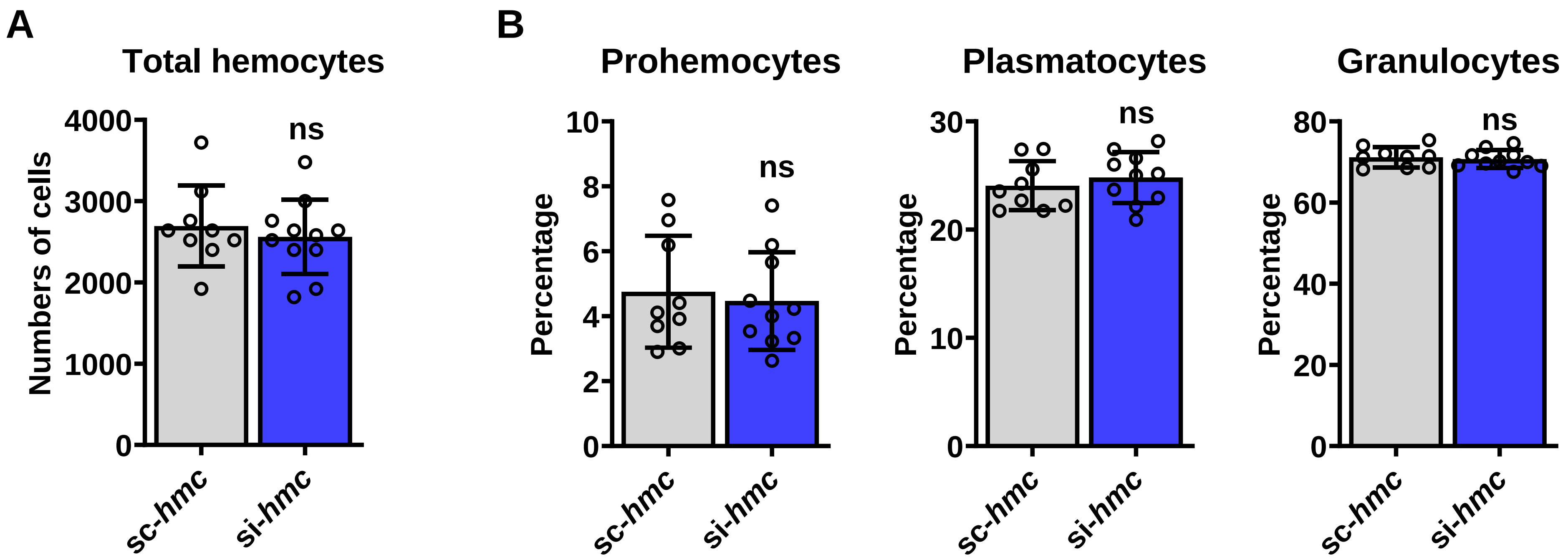

Rolandelli *et al.* - Suppl. Fig. 11

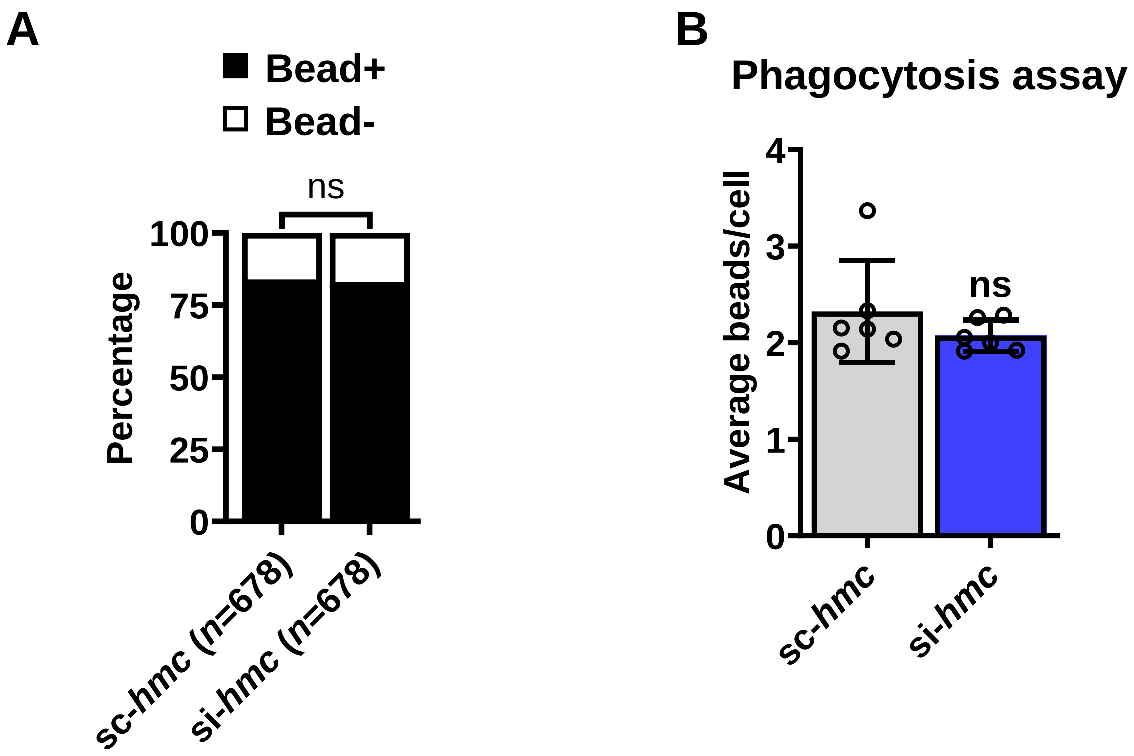

Rolandelli *et al.* - Suppl. Fig. 12

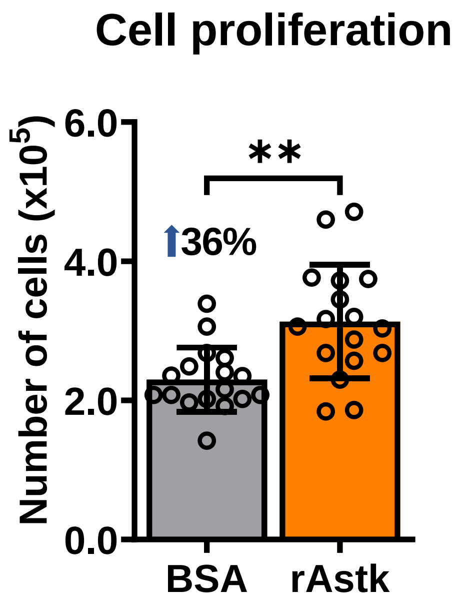

Rolandelli *et al.* - Suppl. Fig. 13

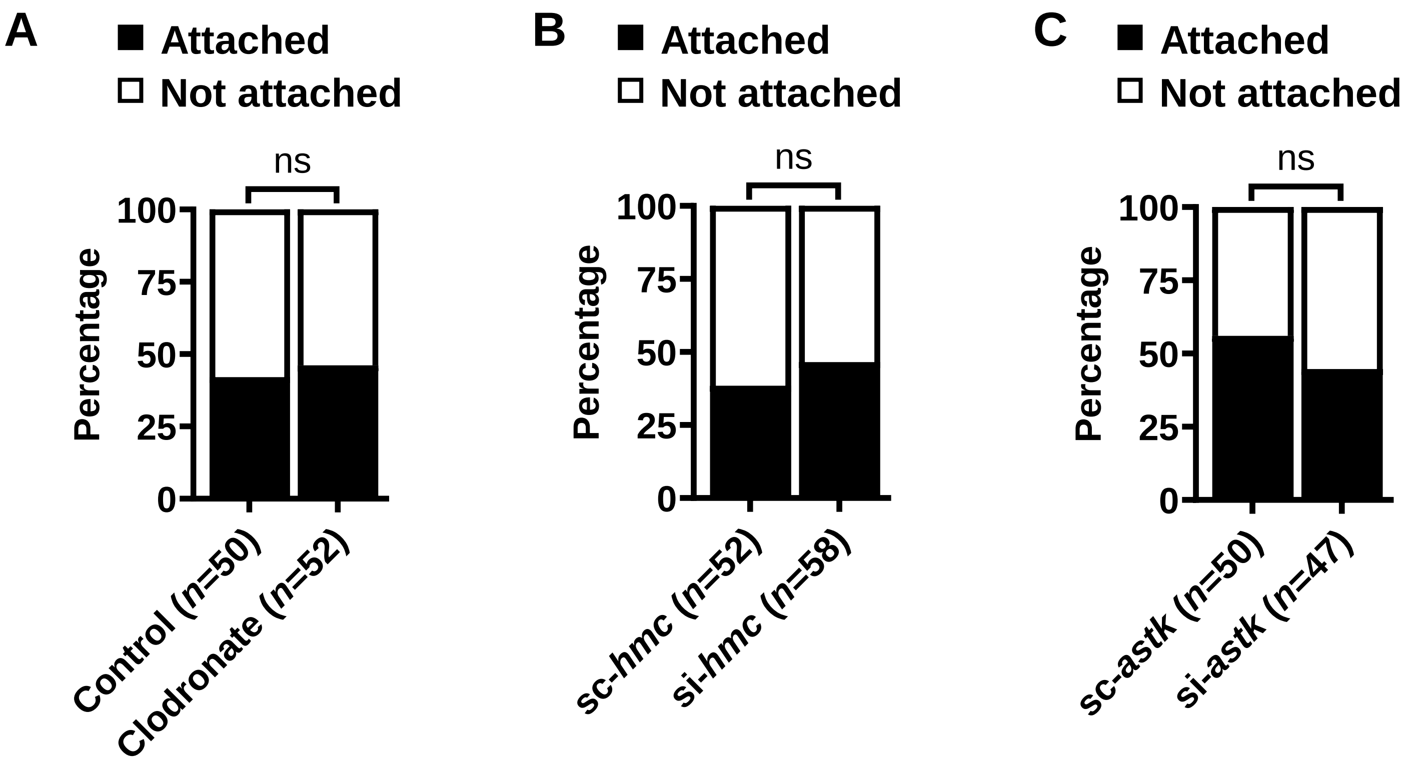

**Rolandelli *et al.* - Suppl. Fig. 14**

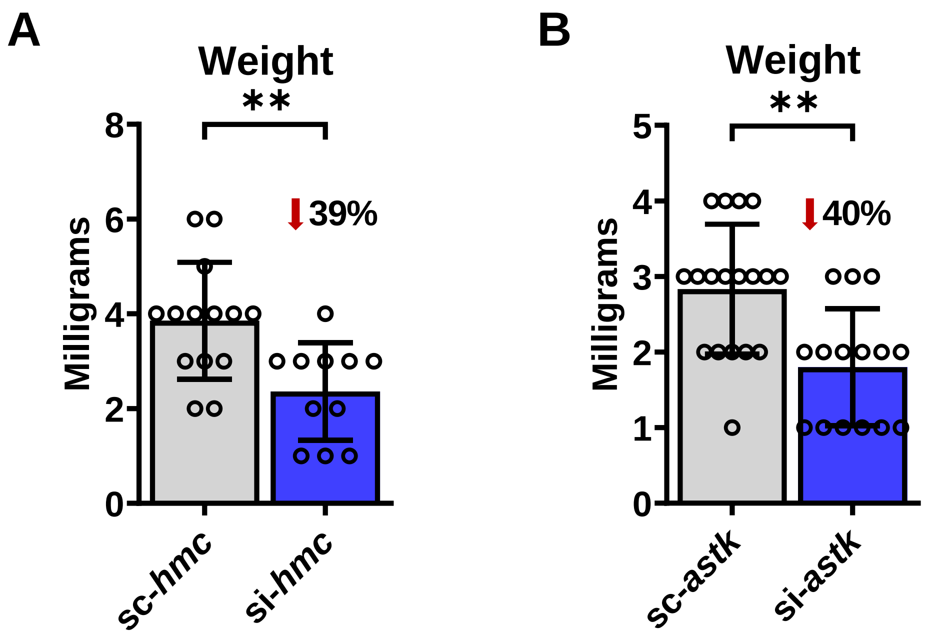

Rolandelli *et al.* - Suppl. Fig. 15

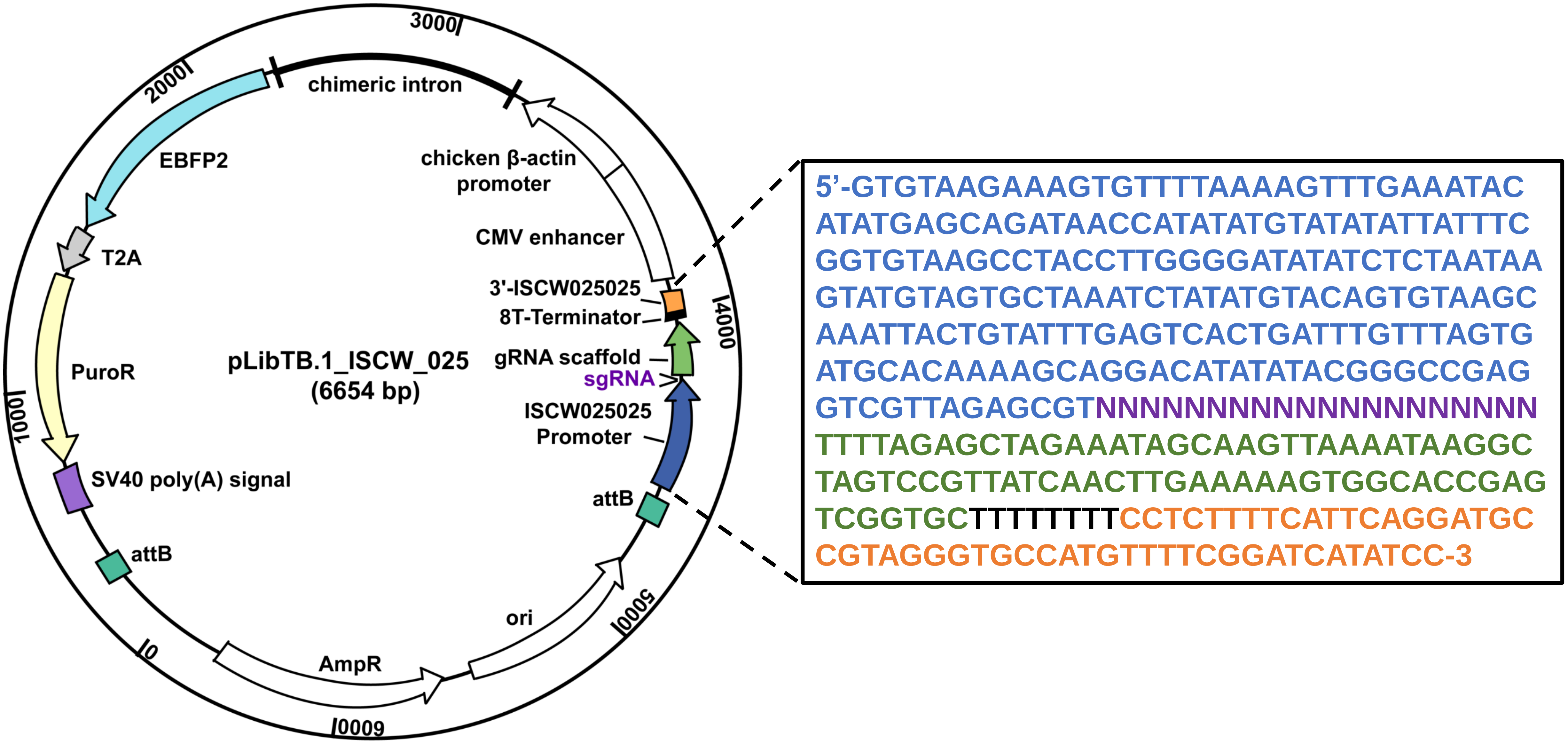

Rolandelli *et al.* - Suppl. Fig. 16
